# Oncogenic mutations in the DNA-binding domain of FOXO1 disrupt folding: quantitative insights from experiments and molecular simulations

**DOI:** 10.1101/2022.04.01.486713

**Authors:** Dylan Novack, Lei Qian, Gwyneth Acker, Vincent A. Voelz, Richard H. G. Baxter

## Abstract

FOXO1, a member of the family of winged-helix motif Forkhead box (FOX) transcription factors, is the most abundantly expressed FOXO member in mature B-cells. Sequencing of diffuse large B-cell lymphoma (DLBCL) tumors and cell lines identified specific mutations in the forkhead domain linked to loss of function. Differential scanning calorimetry and thermal shift assays were used to characterize how eight of these mutations affect the stability of the FOX domain. Mutations L183P and L183R were found to be particularly destabilizing. Electrophoresis mobility shift assays show these same mutations also disrupt FOXO1 binding to their canonical DNA sequences, suggesting the loss of function is due to destabilization of the folded structure. Computational modeling of the effects of mutations on FOXO1 folding was performed using alchemical FEP, and a Markov model of the entire folding reaction was constructed from massively parallel molecular simulations, which predicts folding pathways involving the late folding of helix α3. Although FEP is able to qualitatively predict the destabilization from L183 mutations, we find that a simple hydrophobic transfer model, combined with estimates of unfolded-state solvent accessible surface areas from molecular simulation, is able to more accurately predict changes in folding free energies due to mutations. These results suggest that atomic detail provided from simulation is important for accurate prediction of mutational effects on folding stability. Corresponding disease-associated mutations in other FOX family members support further experimental and computational studies of the folding mechanism of FOX domains.

## Introduction

Forkhead box (FOX) transcription factors are a family of DNA-binding proteins containing a winged-helix motif, a variation of the helix-turn-helix (HTH) motif.^1^ FOX proteins are conserved from *Drosophila* to humans with multiple roles in development and regulation, and mutations in these genes are associated with multiple pathologies including cancer.^2,3^ FOXO1 is a key factor in both insulin signaling and B-cell development, balancing apoptotic and survival signals that vary across tissues and in response to metabolic conditions.^4^

FOXO1 has an N-terminal region, a FOX DNA-binding domain, a nuclear localization signal (NLS), several phosphorylation sites, a nuclear export sequence (NES) and a C-terminal transactivation domain (TAD) (Figure 1A). Nuclear FOXO1 binds to regulatory sites and induces gene expression. Export to the cytoplasm, stimulated by AKT (Ser/Thr protein kinase) phosphorylation leads to ubiquitination and degradation.^5^ The DNA-binding domain has a compact three-helix fold, with the third helix (α3) sitting in the major groove of B-form DNA, and C-terminal β strands projecting along the axis of the DNA to contact one or both of the adjacent minor grooves.^6^

**Figure 1.**
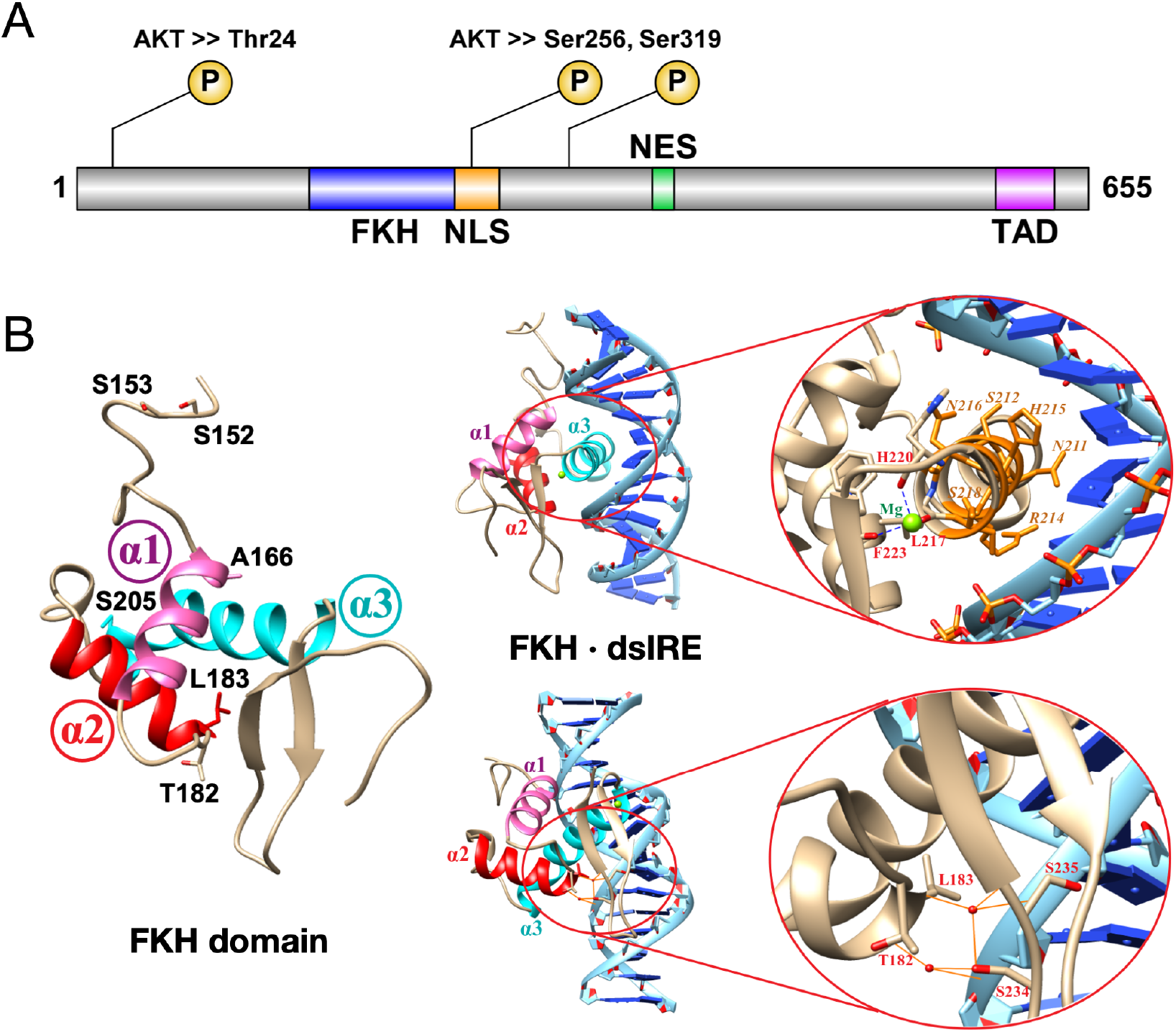
(A) FOXO1 functional domains are shown with three AKT-mediated phosphorylation sites for inactivation. FKH is forkhead DNA-binding domain; NLS is nuclear localization sequence/signal; NES is nuclear export sequence/signal; TAD is transactivation domain. (B) The FKH domain (PDB 3COA) has three helices labeled *α*1 (pink), *α*2 (red) and *α*3 (cyan). The six residues where all eight mutations are found are labeled; none are located in helix *α*3, which makes contacts with binding sequence IRE. Residues T182, L183, S234 and S235 make key hydrogen bonds with the DNA phosphate backbone, mediated by water molecules (red spheres).

FOXO1 is the most abundantly expressed FOXO member in mature B-cells, where it targets genes in apoptosis, cell cycle and growth arrest.^7^ Tonic B-cell receptor (BCR) signaling stimulates the PI3K/AKT pathway via phosphorylated kinase BTK (pBTK) and helps release this brake by cytoplasmic localization of FOXO1. Diffuse large B-cell lymphoma (DLBCL), the most common non-Hodgkin lymphoma among adults,^8^ is fatal if untreated, with ∼40% of patients either unresponsive or relapsing after front-line chemotherapy treatment.^9^ FOXO1 is frequently mutated in DLBCL, and oncogenic driver mutations are associated with cases refractory to, or relapsing after, treatment.^10–12^

Sequencing of DLBCL tumors identified two mutation hotspots in FOXO1, the N-terminus and the FKH domain.^10^ N-terminal mutations in FOXO1 prevent AKT phosphorylation leading to nuclear retention, and are also prevalent and oncogenic in GC-derived tumors such as Burkitt’s Lymphoma.^13,14^ The effects of mutations in the FOX domain, however, have not been investigated. These mutations are spread throughout the folded domain and do not involve residues directly involved in DNA-binding, but mutations at T182 or L183 are adjacent to α3 and the major groove (Fig. 1B).^6,10^ We therefore sought to investigate whether these mutations have a global effect on FKH folding that would disrupt DNA-binding.

To address this question, we performed a joint experimental and computational study of eight oncogenic mutations identified in the FOX domain. Through differential scanning calorimetry and thermal shift assays, we show these mutations destabilize the FOX domain, and in turn disrupt FOXO1 binding to the insulin response element (IRE). Then, as test of state-of-the-art methods to computationally predict the effects of mutations on folding, we compare two simulation-based approaches: (1) an alchemical free energy perturbation (FEP) approach, and (2) Markov state models (MSMs) constructed from massively parallel *ab initio* folding simulations to characterize the folding mechanism of FOXO1. While both approaches reasonably rank-order the effects of destabilizing mutations, we find that changes in per-residue solvent accessible surface area (SASA) extracted from the MSMs, combined with a simple empirical hydrophobicity-based model of protein stability, makes superior quantitative predictions of the

ΔΔ*G* of mutation. These results suggest that detailed information about folding intermediates and unfolded-state structure provided by atomistic simulations is important for accurate prediction of mutational effects on folding stability.

## Results

### Oncogenic mutations affect the stability of the FKH domain

We considered eight FKH mutations identified in DLBCL patients by Trinh et al. (2013),^10^ and two mutations identified in DLBCL cell lines (Figure 1B, Table 1). None of these mutations correspond to residues that directly coordinate DNA bases in canonical DNA recognition sequences such as the insulin response element IRE (Figure 1B, top) or DAF-16 binding elements DBE1 and DBE2.^6^ We therefore hypothesized these mutations may disrupt DNA binding by destabilizing the FKH domain.

**Table 1.** Melting temperatures and enthalpies of unfolding measured by TSA and DSC experiments.

|  | TSA |  | DSC<sup>c</sup> |  |  |  |  |
| --- | --- | --- | --- | --- | --- | --- | --- |
| FKH construct | $T_m$ (°C) | $\Delta T_m$ | $T_m$ (°C) | $\Delta T_m$ | $\Delta H_{cal}$<br>(kcal/mol) | $\Delta H_{vH}$<br>(kcal/mol) | $\Delta\Delta H_{vH-cal}$<br>(kcal/mol) |
| WT | 50.5±0.8 | - | 57.8 ± 0.3 | - | 47 ± 1 | 54 ± 1 | 7 |
| S152R | 51.1±0.1 | 0.6 | 57.0 ± 1.0 | -0.8 | 51 ± 1 | 58 ± 1 | 7 |
| S153R | 51.8±0.1 | 1.2 | 58.6 ± 1.0 | 0.8 | 52 ± 1 | 60 ± 1 | 8 |
| A166G | 46.4±0.5 | -4.2 | 55.1 ± 1.0 | -2.7 | 44 ± 1 | 54 ± 1 | 10 |
| A166V | 56.3±0.9 | 5.8 | 61.9 ± 1.0 | 4.2 | 47 ± 1 | 68 ± 1 | 21 |
| K171E | 52.5±0.8 | 2.0 | – | - | - | - |  |
| T182M<sup>a</sup> | 46.4±0.1 | -4.1 | 51.5 ± 1.0 | -6.3 | 40 ± 1 | 62 ± 1 | 22 |
| L183R<sup>b</sup> | 33.6±1.1 | -16.9 | 41.7 ± 1.0 | -16.1 | 50 ± 1 | 42 ± 1 | -8 |
| L183P | 40.0±0.6 | -10.5 | 48.2 ± 1.0 | -9.6 | 31 ± 1 | 53 ± 1 | 22 |
| S205N | 52.4±1.1 | 1.9 | - | - | - | - |  |
| S205T | 52.7±1.7 | 2.1 | 58.3 ± 1.0 | 0.5 | 58 ± 1 | 59 ± 1 | 1 |

<sup>a</sup>Mutation in OCI-Ly8 cell line. <sup>b</sup>Mutation in WSU-NHL cell line. <sup>c</sup>Wild-type measurements were performed in triplicate (± 0.3 °C), while single measurements were made for mutants (± 1.0 °C)

To test this idea, we performed a thermal shift assay (TSA) for FKH mutants,^15^ using three independent protein batches, three technical replicates per batch (Table 1). According to these assays, the wild-type (wt) FKH domain had melting temperature *T*_m_ = 50.5 ± 0.8 °C. FKH mutants S152R, S153R, K171E, S205N, and S205T had Δ*T*_m_ ≤ 2 °C compared to wt, implying no significant effect on FKH structure. However, significant alterations in *T*_m_ were observed for A166G, A166V, T182M, L183P, and L183R.

In the case of A166, the site of multiple mutations in DLBCL patients, replacement of alanine with the more flexible glycine reduced the melting temperature to 46.4 ± 0.5 °C (Δ*T*_m_ = -4.2 °C) whereas substitution with less flexible valine increases the melting temperature to 56.3 ± 0.5 °C (Δ*T*_m_ = 5.8 °C). However, the most significant mutations were at positions T182 and L183. These residues lie at the N-terminus of helix 2 (*α*2) in the FKH domain. These residues stabilize the overall fold by initiating the first turn of *α*2, and packing against helix 3 (*α*3) which sits in the major groove of bound DNA. When bound to DNA, these residues make water-mediated contacts to the phosphate backbone along with residues 234–235 (Figure 1B).

The mutation T182M—identified in DLBCL cell line OCI Ly8—reduced the melting temperature to 46.4±0.1 °C (Δ*T*_m_ = -4.2 °C) equivalent to A166G. L183P was more destabilizing, reducing the melting temperature to 40.0 ± 0.1 °C (Δ*T*_m_ = -10.5 °C). A second mutation at the same position L183R – identified in DLBCL cell line WSU-NHL – reduced the melting temperature to 33.6 ± 1.1 °C (Δ*T*_m_ = -16.9 °C).

TSA is an indirect measure of protein folding as detection depends on dye binding, which may itself influence the melting temperature Also, the thermal shift assay does not capture the full folding transition as fluorescence above the melting temperature is affected by other non-folding artifacts. Hence we validated our results with an independent measure of protein stability, differential scanning calorimetry (DSC). Two batches of FKH wt and single batches of 8 of the 10 mutants were analyzed using a non two-state folding model to derive melting temperature, the calorimetric and the van’t Hoff enthalpy of unfolding (Table 1).

Melting temperatures determined by DSC were systematically higher by 7 ± 1 °C compared to the thermal shift assay. However, Δ*T*_m_ from DSC closely matched that of the thermal shift assay, with a mean unsigned error (MUE) of 1.3 °C and maximum difference of 2.2 °C (<u>Figure S1</u>). The change in enthalpy (Δ*H*) associated with unfolding were similar between FKH wt and mutants. The wild-type calorimetric enthalpy of unfolding was Δ*H*_cal_ (wt) = 47 ± 1 kcal mol^−1^, while the average value for mutants was Δ*H*_cal_ (mutants) = 47 ± 8 kcal mol^−1^. Similarly, the van’t Hoff enthalpies of unfolding were Δ*H*_vH_ (wt) = 54 ± 1 kcal mol^−1^ vs. Δ*H*_vH_ (mutants) = 57 ± 8 kcal mol^−1^. In general, the van’t Hoff change in enthalpy was higher than the calorimetric enthalpy change by 10 ± 10 kcal mol^−1^, suggesting that reductions in melting temperature of FKH mutants result from perturbations to the folded or unfolded states, but not a gross change in the folding landscape of the FKH domain.

### Mutations that reduce FKH stability also reduce DNA binding

We expected that FKH mutations identified in DLBCL cell lines and patient specimens which significantly destabilize the FKH domain should also reduce binding to canonical DNA binding sequences such as the insulin response element (IRE) and Daf-16 binding element (DBE). We therefore performed electrophoresis mobility shift assay (EMSA) experiments to measure the binding of FKH wt, L183P and L183R to IRE and DBE2 oligonucleotides (Table 2). Experiments were performed using three independent protein batches, and bands quantified by ImageLab (BioRad). The results were fit to a two-state binding model to estimate *K*_D_.

**Table 2.**
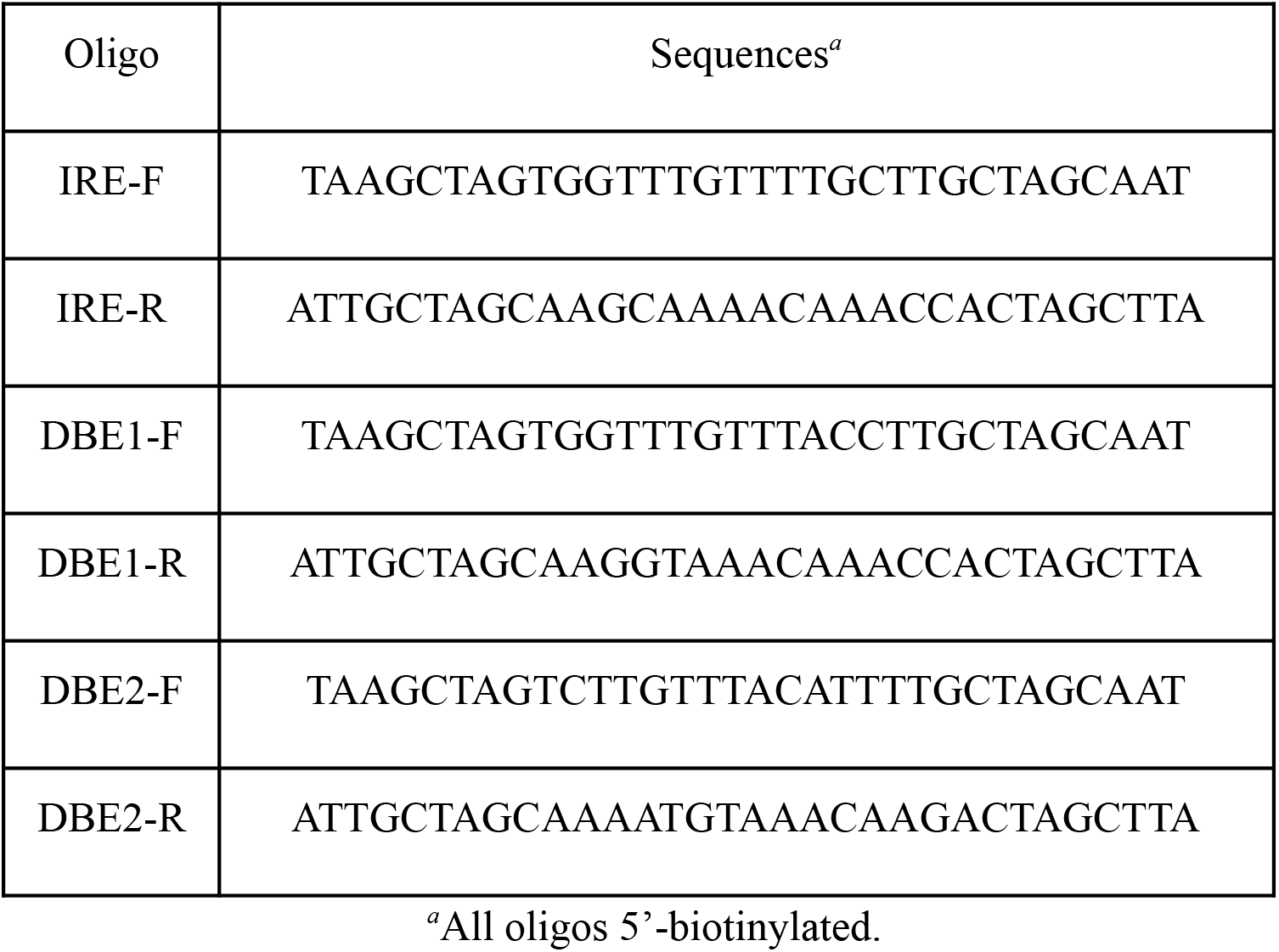
DNA oligonucleotide sequences used for FKH binding studies.

Representative EMSA images clearly illustrate that L183P and L183R show progressively lower affinity for dsDNA (Figure 2). The affinity of FKH wt for IRE from three independent experiments was *K*_D_ = 200 ± 70 nM, ∼2-fold weaker than previously reported.^25^ The affinity of FKH L183P for IRE was 2.5-fold weaker, *K*_D_ = 460 ± 100 nM. The affinity for L183R for IRE was significantly weaker, the dissociation constant was not measurable for the concentration tested, *K*_D_ > 1 µM.

**Figure 2.**
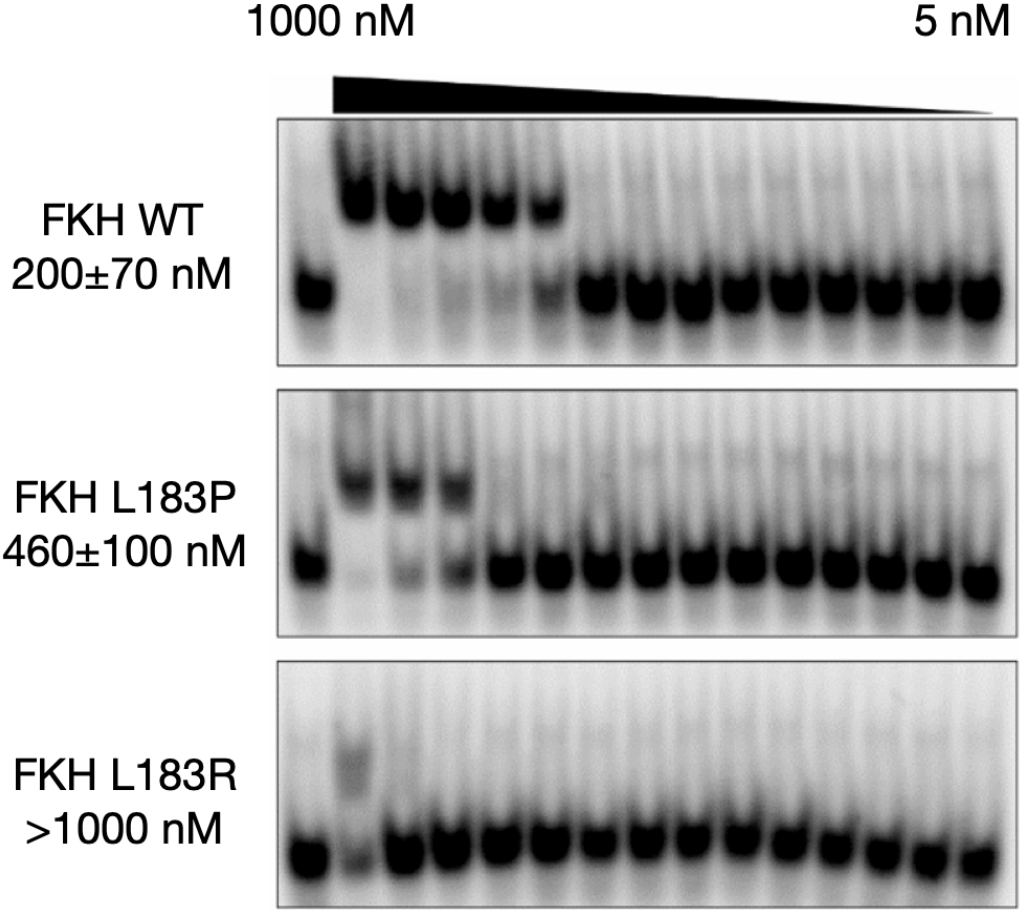
EMSA analysis of FKH binding to the insulin response element (IRE). Representative blots of FKH wt, FKH L183P and FKH L183R serial dilution from 1 µM (1000 nM) to 5 nM in complex with 1 nM dsIRE. Each blot is labeled with *K*_D_ estimates in nM.

The EMSA assay was repeated for dsDBE2 with single batches of FKH wt and FKH L183P. The affinity of FKH wt for DBE2 was *K*_D_ = 90 nM, 10-fold weaker than previously reported, but again the affinity of FKH L183P was 2.5-fold weaker, *K*_D_ = 230 nM. These results support the conclusion that FKH mutations which disrupt folding also disrupt binding to canonical target DNA sequences.

### FEP calculations partially explain how FKH mutations disrupt folding

We next sought to more fully characterize structural mechanisms by which mutations destabilize the FOXO1 FKH domain, using alchemical free energy perturbation (FEP, see <u>Materials and Methods</u>). This method relies on atomistic molecular dynamics simulation to calculate the free energy cost of transforming a wild-type amino acid residue to a mutant one.^16^ The difference between this free energy cost for the folded state (calculated using the folded FKH domain), and the free energy cost for the unfolded state (calculated using a tripeptide model of the unfolded state), yields a prediction of the change in the folding free energy upon mutation, ΔΔ*G*_fold_ (Figure 3A). The accuracy of this state-of-the-art method is expected to be 1.1–1.6 kcal mol^−1^ (MUE) based on a number of recent benchmarks.^17–19^

**Figure 3.**
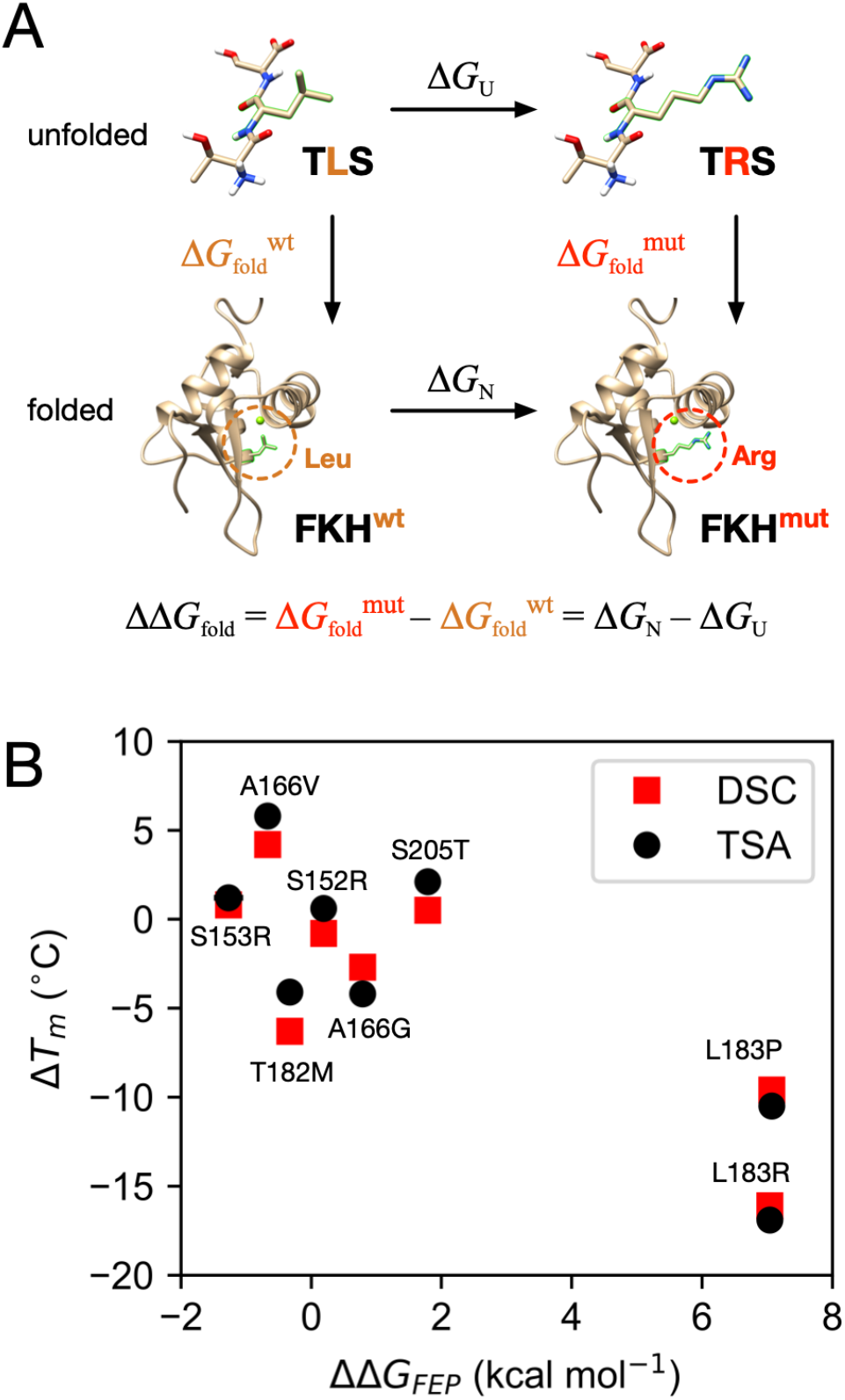
(A) A thermodynamic cycle illustrates how FEP is used to calculate ΔΔ*G*_fold_ for eight FOXO1 FKH mutants, with the L183R mutant shown as an example. (B) Correlation of ΔΔ*G*_fold_ values calculated using FEP (ΔΔ*G*_FEP_) with Δ*T*_m_ from TSA (black) and DSC (red). Uncertainty estimates for ΔΔ*G*_FEP_ are smaller than the marker width.

The FEP results qualitatively agree with the experimental Δ*T*_m_ measurements (Pearson correlation coefficient of *r* = -0.85 for both TSA and DSC measurements, Figure 3B). Most importantly, FEP predicts the most destabilizing mutations to be L183P and L183R, in agreement with experiments, but for mutations that change the melting temperature by less than ±5 °C, FEP often fails to predict over-vs. under-stabilization, particularly for FKH T182M and S205T.

To quantitatively evaluate the accuracy of the FEP predictions, we used the the DSC *T*_*m*_ values for each for each FKH variant to estimate experimental ΔΔ*G*_fold_ using a method adapted from Robertson, Murphy, and Rees (see Materials and Methods).^20–22^ This comparison reveals that poor quantitative agreement between FEP estimates of ΔΔ*G*_fold_ and experiment, especially for mutations that significantly decrease stability of the FOXO1 FKH domain (see Figure 7C), with a MUE of 2.11 kcal mol^−1^ (RMSE 3.5 kcal mol^−1^). Aside from protein force field issues (which have been found to be generally robust for relative FEP ^18^), there are several reasons why this may be. One possibility is that the unfolded state is poorly approximated by the tripeptide simulation; by comparison, realistic protein unfolded states may be significantly more compact, with specific residual structure.^23–25^ Another reason may stem from the well-known challenges associated with charge-changing and proline transformations.^18,26^

### Massively parallel simulations and Markov models elucidate the folding mechanism of FOXO1 at atomic resolution

To better understand the folding of FOXO1 FKH domain at higher-resolution, and how oncogenic mutations might perturb this process, we next sought to construct a Markov model of the folding mechanism,^27–31^ from nearly ten thousand independent molecular simulation trajectories simulated on the Folding@home distributed computing platform.^32,33^ Unlike previous *ab initio* folding studies, which have focused on well-studied mini-proteins,^34,35^ the folding of FOXO1 has not yet been experimentally characterized.

Simulation trajectories were started from twenty different folded and unfolded conformations generated from high-temperature simulations (<u>Figure S2</u>, see Materials and Methods), with production runs performed at 375 K, which we estimated to be close to the simulation melting temperature of FOXO1 FKH based on previous work for proteins of similar size and topology.^36^ An aggregate ∼6.8 ms of simulation data was collected from 9927 trajectories generated using GPU-accelerated OpenMM ^37^ with the AMBER ff14SB force field ^38^ and TIP3P solvent.^39^

Time-lagged independent component analysis (TICA) was used to project the trajectory data to components that best capture the slowest motions in the folding reaction.^40,41^ Trajectories were featurized using pairwise distances between every other backbone C_α_, and the time-lagged correlation matrix of these features was computed using lag time of 2.5 ns. Projection of the trajectory data onto the first two time-independent coordinates tIC1 and tIC2 shows a well-sampled funnel-shaped landscape, with the slowest motions along tIC1 corresponding to global folding/unfolding (Figure 4A).

**Figure 4.**
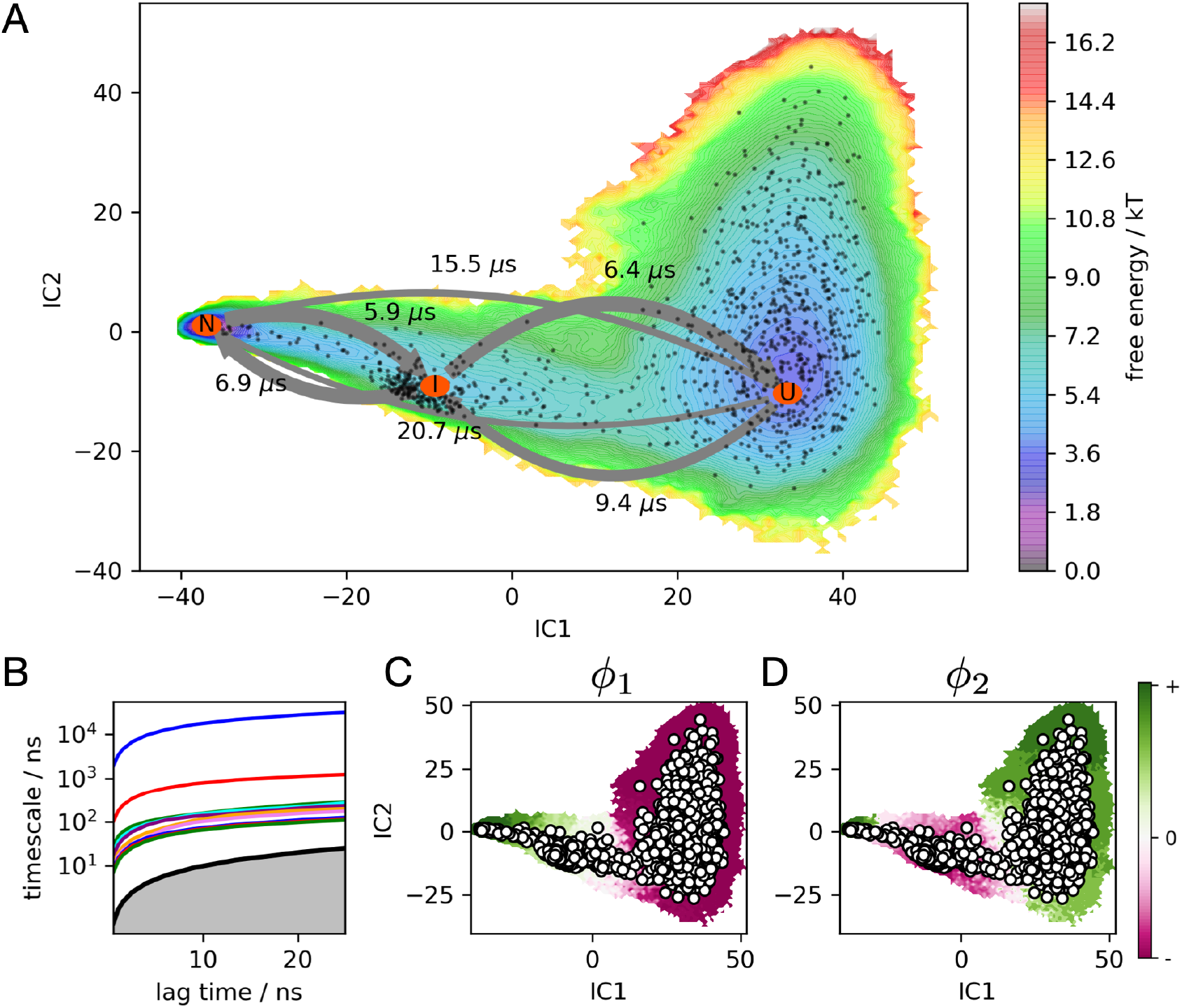
(A) FOXO1 folding free energy landscape at 375 K calculated from the MSM, projected to the two largest TICA components. MSM macrostates U, I, and N are labeled with orange ovals, connected by arrows whose widths represent transition frequencies, labeled with mean first passage times. Black dots represent MSM microstates. (B) An implied timescale plot shows the ten slowest timescales as a function of the MSM lag time. (C and D) The first and second relaxation eigenmodes from the MSM, colored by sign structure. The first eigenmode shows population flux from N to U at 375 K, while the second eigenmode shows flux into I from U and N.

Trajectory data projected to the four largest TICA components were clustered into 1000 states using a *k*-centers algorithm, enabling the construction of a Bayesian MSM, a method which reduces the statistical error of model estimation.^42^ The first relaxation eigenmode of this model shows population flowing from folded (+) to unfolded (-) conformations along tIC1, indicating that the protein is moderately unstable at the simulation temperature of 375 K. Although unfolding events are observed more frequently than folding events, many folding and refolding events are also observed (<u>Figure S3</u> and <u>Figure S4</u>). Using PCCA+^43,44^ we clustered the individual microstates into three coarse-grained macrostates identified as unfolded (U), intermediate (I), and folded (N) (Figure 5). Mean passage times between macrostates are on the 5–20 µs timescale (<u>Table S1</u>), with the mean first passage time of folding estimated to be 20.7 *µ*s.

**Figure 5.**
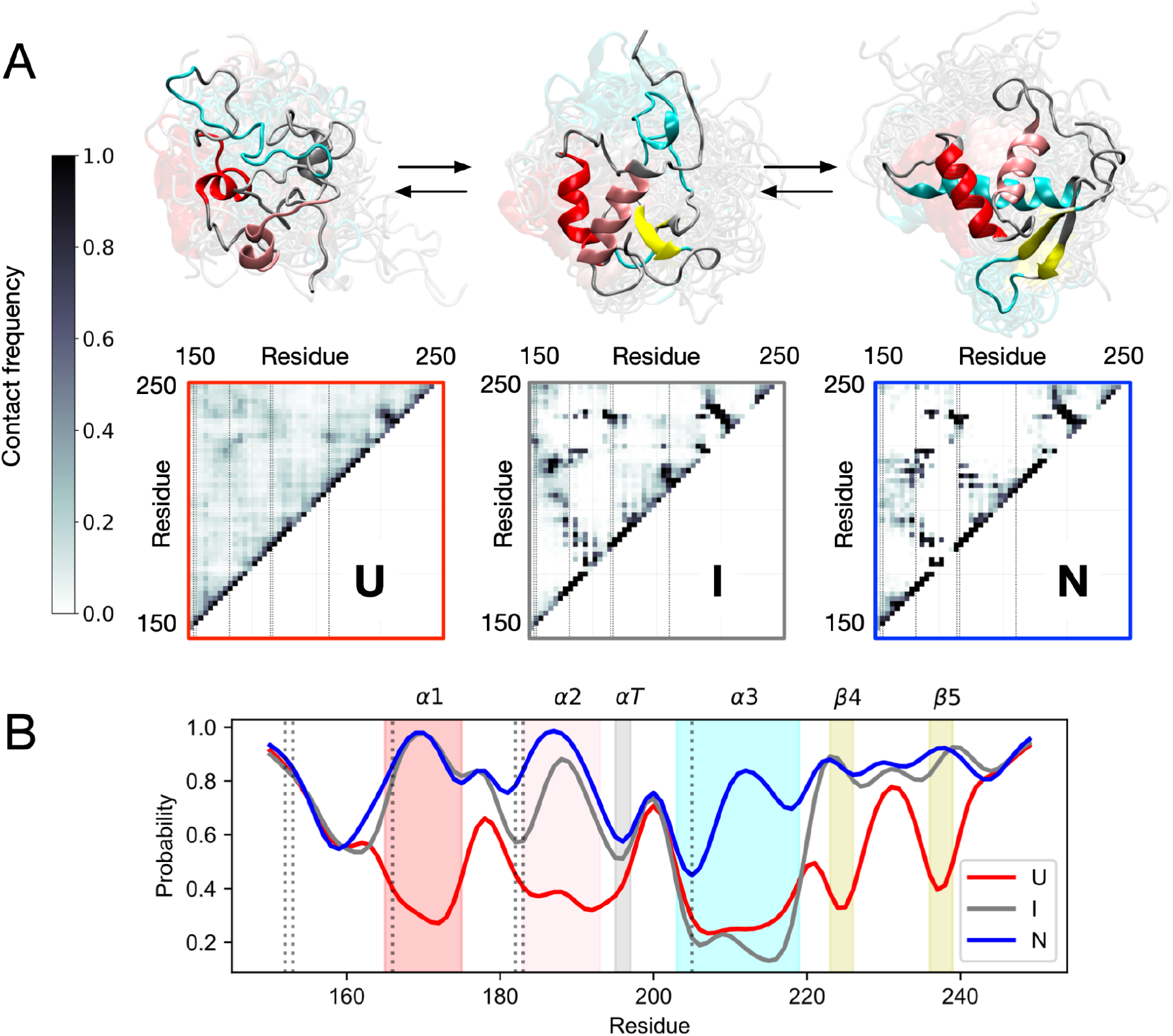
Simulations predict helix α3 formation to be the final step in folding. (A) Interresidue contact frequencies and representative structures of MSM macrostates U, I and N. Vertical lines on the contact maps denote mutant positions. (B) Per-residue populations of native secondary structure for each macrostate. Shaded bands mark native secondary structures α1, α2, αT, α3, β4 and β5, with vertical lines denoting mutant positions.

To elucidate the main structural events in the folding pathway, the DSSP algorithm^45^ was used to assign per-residue secondary structure to each macrostate and the populations of native secondary structure elements were calculated (Figure 5A). FOXO1 has three *α*-helices (*α*1, *α*2, *α*3), an *α*-turn (*α*T), and two *β*-sheets (*β*4, *β*5). In the unfolded state (**U**), the probability *p* of any of these structures to be correctly folded is less than 0.5. In the intermediate state (**I**), we find that there is a high probability (*p* > 0.8) that secondary structures are formed except for *α*3 (0.2 ≤ *p* ≤ 0.3). Only after the other structural elements are in place does *α*3 form in the native state (**N**). An analysis of interresidue contact frequencies for each macrostate reaches similar conclusions (Figure 5B).^46^ Native contacts have low populations in **U**, but are mostly formed in **I** except for contacts with *α*3. In **N** all native contacts are formed. This is consistent with sequence-based predictions from PSIPRED of poor helix propensities for *α*3 (<u>Figure S2</u>), suggesting tertiary context is important for its formation.^47^ This context-dependence may explain why none of the highly destabilizing mutations are located in *α*3, except S205, which is far from the major groove when bound to DNA.^6^

While the crystal structure of DNA-bound FOXO1 contains a Ca^2+^ ion that helps stabilize *α*3, the context-dependent folding of *α*3 is likely a general feature of FOXO domain structure and dynamics. NMR structural studies of four different FOXO domains in the absence of Ca^2+^ show differences in solution-state structure at *α*T and *α*3, but highly similar crystal structures when bound to DNA.^48^

### A hydrophobic transfer model accurately predicts thermostability changes of mutations from simulated changes in SASA

Simulations predict a compact denatured state for FOXO1 with populations of residual structure. We hypothesized that the structural detail and extensive statistical sampling provided by simulations could help us evaluate whether the tripeptide model used in the FEP studies is a sufficiently accurate approximation of the unfolded state.

The Shrake-Rupley algorithm^49^ was used to calculate how the average solvent-accessible surface area (SASA) of residues S152, S153, A166, T182, L183, and S205 changes for each macrostate along the folding reaction (Figure 6). Interestingly, predicted changes in SASA were highly non-trivial. In many cases, the SASA of the native state (N) was comparable (S152 and S153) or greater (S205) than the unfolded state (U). More importantly, for all residues, the average SASA in the tripeptide model of the unfolded state used for FEP was at least 2.8 times larger than in our simulations, suggesting that a more realistic model of SASA in the unfolded state might lead to more accurate predictions of how mutations change FOXO1 stability.

**Figure 6.**
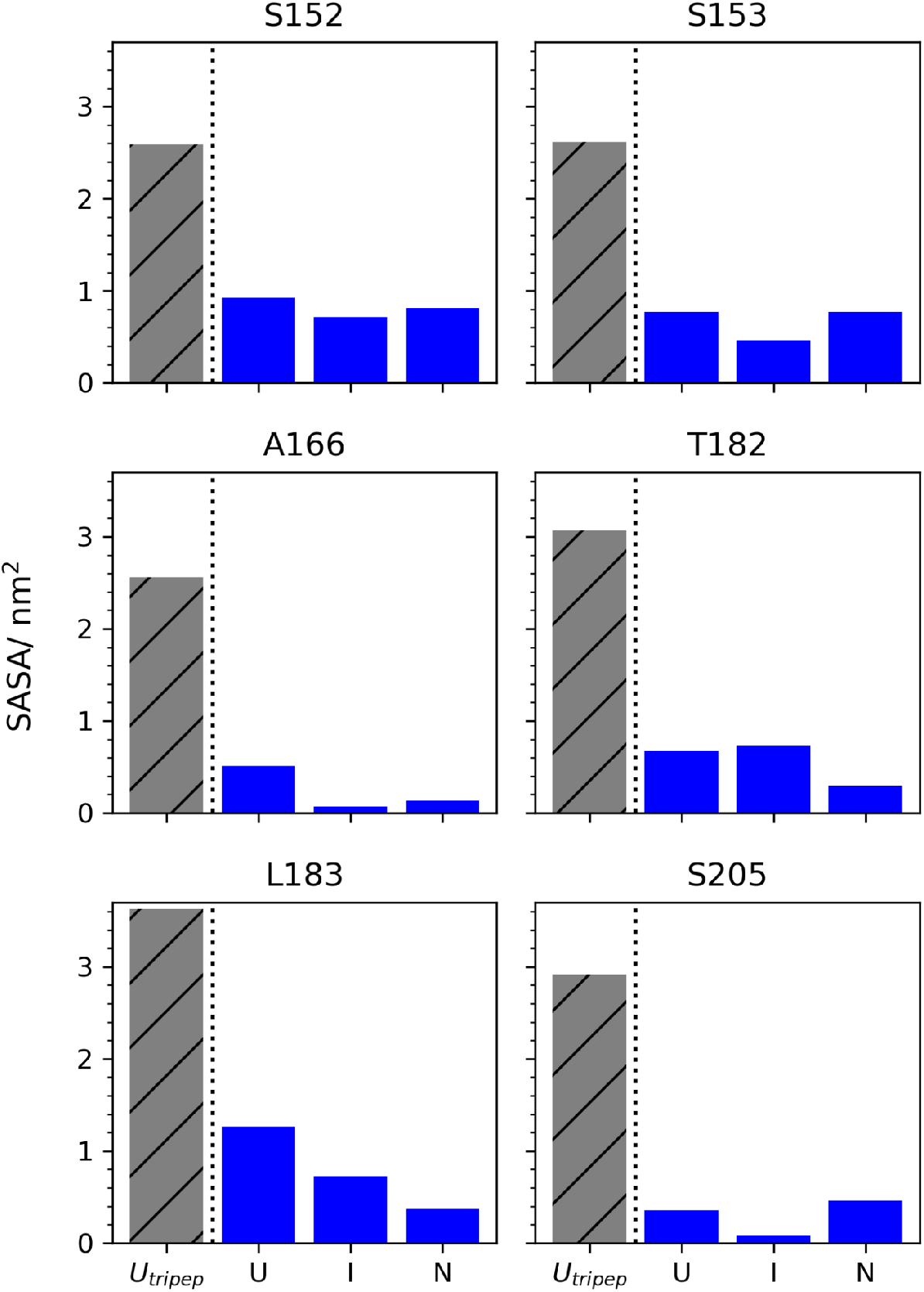
Tripeptide models of the unfolded state consistently overestimate per-residue solvent-accessible surface area (SASAs) compared to molecular simulations. For each mutant position (S152, S153, A166, T182, L183 and S205), bar graphs show average SASA calculated from tripeptide FEP simulations (patterned gray), versus average SASA of MSM macrostates U, I, and N (blue).

To test this idea, we constructed a model of how the free energy of folding Δ*G* (U→N) depends on changes in the local hydrophobic environment, based on the empirical hydrophobic transfer model of Eisenberg et al.^50–53^ The change in the free energy of folding ΔΔ*G*_fold_ due to a mutation at residue position *i*, from amino acid residue *r*_*i*_ to *s*_*i*_, is computed as

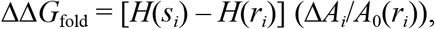

where *H*(*s*_*i*_) and *H*(*r*_*i*_) are the hydrophobicities of the mutant residue *s*_*i*_ and wild type residue *r*_*i*_, respectively, Δ*A*_*i*_ is the predicted change in SASA of (the wild type) residue *i* upon folding (U→N), and *A*_0_(*r*_*i*_) is the maximum solvent exposure of the amino acid residue at position *i*. Importantly, the only free parameter in this model is the input value Δ*A*_*i*_, which we compute directly from the molecular simulations (see <u>Materials and Methods</u>). All other parameters come directly from previously published work: the hydrophobicities *H* come from the consensus hydrophobicity scale of Eisenberg et al (1982), ^51^ and the normalized maximum solvent exposures of amino acids *A*^0^_*i*_ are taken from Tien et al.^53^

For comparison, we also calculated ΔΔ*G*_fold_ estimates using the popular FoldX algorithm, a native structure-based empirical predictor of protein stability trained on a large corpus experimental data.^54^

Figure 7 shows a side-by-side comparison of three different computational predictions of ΔΔ*G*_fold_: from the hydrophobic transfer model (Figure 7A), from FoldX (Figure 7B), and from FEP calculations (Figure 7C). Of these, the hydrophobic transfer model agrees most accurately with the DSC results, with a RMSE of 0.25 kcal mol^−1^, and a linear fit of slope 0.98. The FoldX predictions generally agree with results from DSC, but overestimate the magnitude of destabilizing mutations (RMSE = 0.87 kcal mol^−1^). The FEP predictions are least accurate, severely overestimating the magnitude of destabilizing mutations with RMSE = 3.1 kcal mol^−1^ and a linear fit slope of 5.09.

Interestingly, all three methods predict T182M to be stabilizing, despite the experimental finding that it is destabilizing (ΔΔ*G*_fold_ = +0.42 kcal mol^−1^). This may be due the the context-dependent importance of threonine at that position in the native structure, where it serves both as a strand-pairing residue in a preferred beta-sheet conformation, and as a N-terminal capping group, whose side chain makes a backbone hydrogen bond to stabilize *α*2.

**Figure 7.**
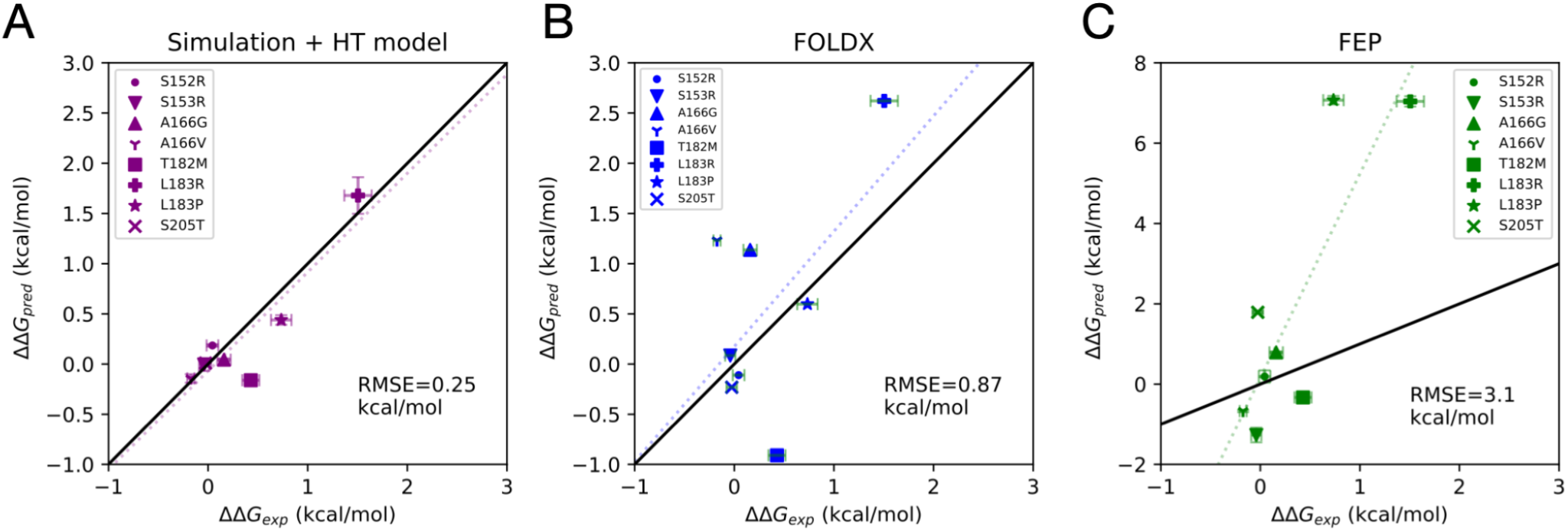
Comparison of estimated experimental folding free energy changes upon mutation ΔΔ*G*_exp_, and predicted estimates ΔΔ*G*_pred_ from three different computational methods: (A) a combined molecular simulation and hydrophobic transfer (HT) model, (B) predictions from the FoldX algorithm, and (C) alchemical FEP estimation. In each plot, the dotted line is the least-squares linear fit, and the root mean squared error (RMSE)

### FEP calculations along the folding coordinate show a continuum of disruption to local folding

To further explore the mechanism by which mutations perturb the folding reaction, six representative structures of folding intermediates were taken along the folding coordinate tIC1, and used as fixed-backbone structural templates for FEP calculations of the free energy of mutation (Figure 8A). For most of these structures, helices *α*1 and *α*2 are folded or partially folded.

**Figure 8.**
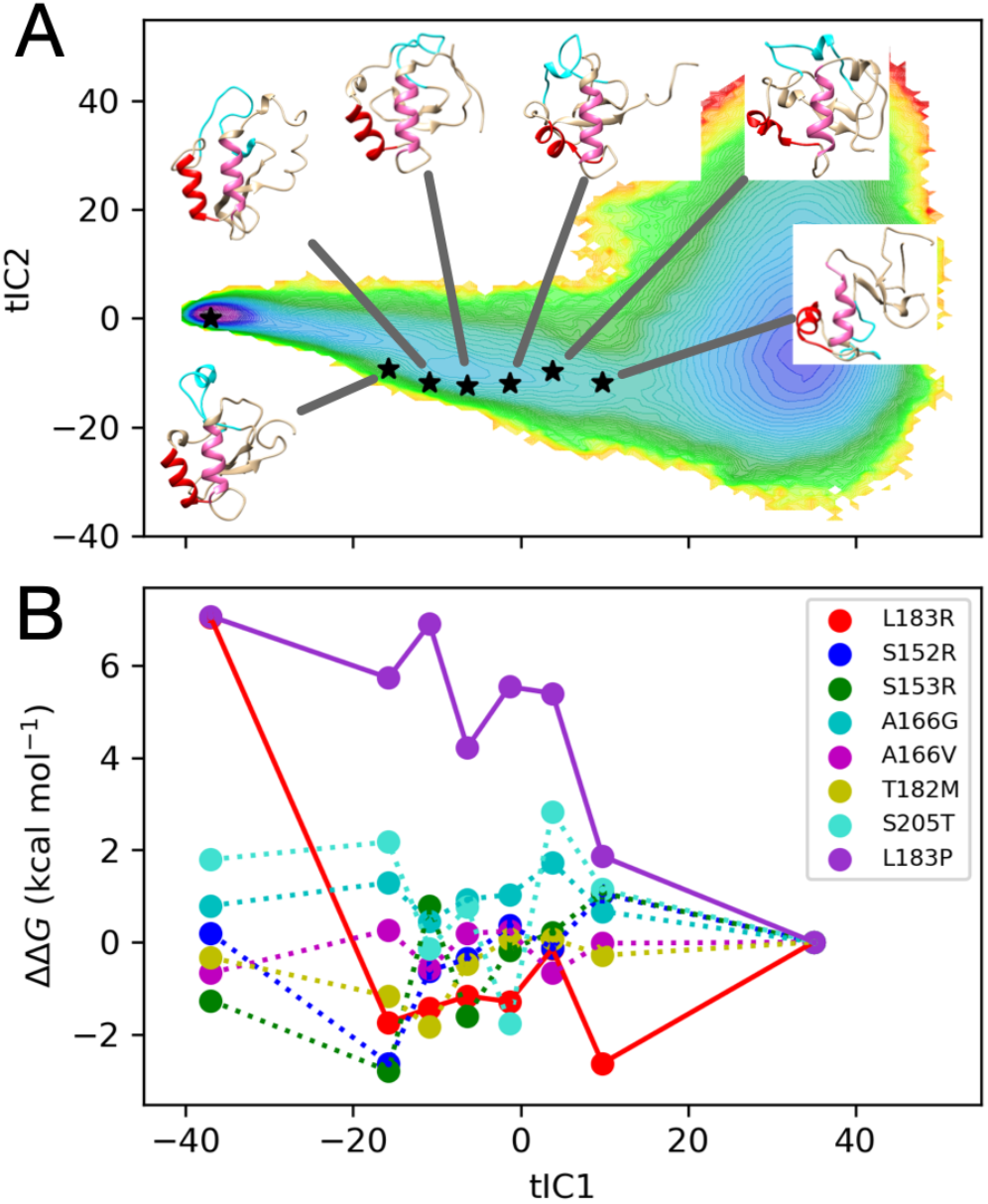
(A) Six folding intermediates *i* along the tIC1 folding coordinate were chosen as templates for FEP estimation of ΔΔ*G* = Δ*G*_*i*_ – Δ*G*_U_, where the tripeptide model of the unfolded state U is used as a reference. (B) Estimates of ΔΔ*G* along tIC1 suggest that perturbations from mutations are dependent on the folding reaction coordinate, particularly for L183R and L183P (solid lines).

Figure 8B shows a plot of the predicted ΔΔ*G* = Δ*G*_*i*_ – Δ*G*_U_ from FEP, where Δ*G*_*i*_ is calculated using one of the structures along the folding coordinate tIC1, and Δ*G*_U_ is taken from the tripeptide model of the unfolded-state. When Δ*G*_*i*_ = Δ*G*_N_, ΔΔ*G* recapitulates the FEP predictions of ΔΔ*G*_fold_, but for intermediate Δ*G*_*i*_, the value of ΔΔG reports on the magnitude of the perturbation along the folding reaction. The results suggest that destabilizing mutations L183P and L183R may exert their effects differently along the folding reaction: L183P perturbations gradually increase as FOXO1 acquires native structure, while L183R appears to particularly destabilize the **N** state in the last step of folding.

## Discussion

To the best of our knowledge, this work represents the first detailed investigation of FOX domain folding mechanism. The Markov model we have constructed makes testable predictions about the folding pathways and rates of this ‘winged helix’ motif that can be tested experimentally. Previous studies of Engrailed homeodomain (EnHD) and its homolog Pit1 demonstrate the malleable stability of the HTH motif, i.e. α2-α3 of the FOX domain, resulting in a continuum of folding mechanisms from ultrafast three-state framework to apparent two-state nucleation-condensation model.^55^ We speculate that similar malleability might be seen across the family of FOX domains, an intriguing area to explore in the future within the context of disease-related mutations.

A key result of this study is that two oncogenic point mutations at L183 dramatically destabilize the FOX domain, resulting in a FKH domain that is largely unfolded at physiological temperatures, and loss of function (DNA-binding).

Point mutations that destabilize the FOX domain have been previously reported. Mutation of paired residues within the β-turn ‘wing1’ of FOXO1 homolog FOXD3 to glycine and proline reduced the *T*_m_ by 3 °C and 5 °C, respectively.^56^ A naturally occurring mutation in FOXD3, N173H associated with developmental ocular conditions, corresponds to FOXO1 K192; this occurs on the protein surface and is not expected to affect folding or DNA binding.^57^ Two mutations in FOXC1, L130F and W152G also associated with developmental ocular conditions, correspond to conserved residues FOXO1 L217 and W237.^58,59^ FOXO1 L217 lies on helix α3 and W237 on the second β-strand of ‘wing1’, both within 4 Å of L183. Assays of both FOXC1 mutations in cell culture indicated reduced DNA-binding and formation of protein aggregates in the cytoplasm;^58,59^ cytoplasmic localization of FOXO1 is also observed in DLBCL specimens.^60,61^ Thermal analysis of six disease-causing mutants of FOXG1 all showed dramatic reductions in *T*_m_, with Δ*T*_m_ ranging from –8 to –15 °C relative to wild-type. These included FOXG1 R230, corresponding to conserved FOXO1 R214 that—like L217—lies on helix α3 within 4 Å of L183.^62^ Taken together, there are three different residues with 4 Å of FOXO1 L183 with either in vivo and or vitro evidence of disrupted folding, suggesting a common conserved hydrophobic core involving the packing of helices α2 and α3. The loci of native-state contacts with these residues supports this idea (Table 4).

**Table 3.** Estimated ΔΔ*G*_fold_ values for FOXO1 from DSC melting temperatures, and predicted ΔΔ*G*_fold_ values from three different computational methods: a hydrophobic transfer model (ΔΔ*G*_HT_), the FoldX algorithm (ΔΔ*G*_FoldX_), and free energy perturbation (FEP, ΔΔ*G*_FEP_). All values are reported in kcal/mol.

| Mutation | $\Delta\Delta G_{\text{fold}}$ (DSC)<sup>a</sup> | $\Delta\Delta G_{\text{HT}}$ | $\Delta\Delta G_{\text{FoldX}}$ | $\Delta\Delta G_{\text{FEP}}$ |
| --- | --- | --- | --- | --- |
| S152R | $0.04 \pm 0.06$ | $0.185 \pm 0.01$ | -0.110 | $0.19 \pm 0.15$ |
| S153R | $-0.04 \pm 0.05$ | $-0.004 \pm 0.06$ | 0.085 | $-1.27 \pm 0.18$ |
| A166G | $0.16 \pm 0.07$ | $0.044 \pm 0.01$ | 1.137 | $0.79 \pm 0.02$ |
| A166V | $-0.17 \pm 0.04$ | $-0.143 \pm 0.04$ | 1.231 | $-0.67 \pm 0.03$ |
| T182M | $0.43 \pm 0.09$ | $-0.161 \pm 0.03$ | -0.911 | $-0.33 \pm 0.06$ |
| L183R | $1.51 \pm 0.14$ | $1.679 \pm 0.18$ | 2.621 | $7.04 \pm 0.13$ |
| L183P | $0.73 \pm 0.10$ | $0.44 \pm 0.05$ | 0.597 | $7.07 \pm 0.11$ |
| S205T | $-0.03 \pm 0.05$ | $0.009 \pm 0.002$ | -0.231 | $1.79 \pm 0.03$ |

<sup>a</sup>calculated from the theory of Robertson and Murphy, using $N_{\text{res}}=100$ (see Materials and Methods).

**Table 4.** The location of native-state contacts for residues with selected oncogenic mutations.

| **Residue** | **Crystal Contacts** |
| --- | --- |
| A166 | T170, N216, H220, F223 |
| T182 | I186, N228, S234, S235, M236, W237 |
| L183 | T187, I213, R214, L217, S234, S235, W237 |
| S205 | D199, K200, G201, W209 |

The destabilization of the FOXO1 fold by L183P and L183R has important translational significance. The loss of stability abrogates DNA-binding, implying loss-of-function (LOF) and oncogenicity due to the negation of canonical tumor suppressor function of FOXO1 in mature B cells. We note that: (i) although FOXO1 is one of the more commonly mutated genes in DLBCL, L183 mutations are overall extremely rare, (ii) cancer mutations are heterozygous, there should be a second wild-type allele to maintain FOXO1 function, and (iii) most of the 10 FOXO1 mutations tested had little to no effect on folding of the FOX domain. There are many other ways mutations in the FOX domain could affect FOXO1 function, and further cell-based assays are required. Nevertheless, we also note that FOXO1 cytoplasmic relocalization is relatively common in DLBCL cases, just as BCL2/MYC overexpression is more common than genetic translocation.^61^ FOXO1 L183 mutation could be a genetic instance of a more general phenotype of unfolding/misfolding of FOX domains, and this could result not only in LOF phenotypes but toxic gain-of-function (GOF) such as off-target or nonspecific DNA-binding and gene transcription. These observations, together with corresponding disease-associated mutations such as in FOXC1 and FOXG1,^63,64^ support further experimental and computational studies of the folding mechanism of FOX domains.

To better understand how disease-related mutations destabilize FOXO1, we performed both FEP calculations and massively parallel molecular simulations of the folding reaction. Were these calculations worth the expense? Yes, for several reasons. First, the destabilizing effect of L183P is not obvious from inspection of the native state alone; proline is not uncommon in the first turn of alpha helices, in fact L183 corresponds to the position of highest propensity (Ncap+1).^65^ However, the restricted range of dihedral angles for proline clearly disrupt the folding of the HTH motif. Similarly, the destabilizing effect of arginine is not obvious from examination of the native state; the FOX domain is already highly basic with 8 arginines and 12 lysines in 115 residues facilitating favorable electrostatic interaction with the DNA backbone. However, the introduction of a bulky charged residue into the hydrophobic core of the HTH motif clearly destabilizes the HTH motif and hence the folding of the FOX domain. These observations of L183, together with the milder effect of FOXO1 T182M and identification of corresponding destabilizing mutations in FOXC1 and FOXG1, emphasize the value of modeling the folding landscape of FOXO1 in atomic detail.

Another reason to perform atomistic folding simulations is to accurately model the unfolded state. While we found FEP predictions of the destabilizing effects of mutations to be qualitatively instructive, the hydrophobic transfer model is able to infer accurate *quantitative* predictions from simulated unfolded states. One likely reason for the inaccuracy of FEP is the use of an unrealistic tripeptide model, which, as we have shown, severely overestimates the extent of solvent exposure in the unfolded state. On the other hand, our FEP studies along the folding coordinate (Figure 8) show that even if we assume that the unfolded state of FOXO1 resembles a compact and partially folded intermediate, FEP still overestimates the magnitude of destabilization.

While FEP remains state-of-the-art in terms of molecular modeling, “exaggerated” predictions of the effects of mutations are not uncommon. Gapsys et. al. (2016) analyzed 119 mutations in the enzyme barnase by alchemical FEP, and found the the range of calculated global ΔΔ*G*_fold_ to be -2.2–7.2 kcal mol^−1^.^66^ Kucukkal et al., studying the effects of 10 Rett Syndrome mutations in MeCP2 MBD, note that accurately predicting the effects of charge changing mutations via FEP is particularly difficult.^67^ Steinbrecher et al., in a large-scale validation of FEP+, provide additional evidence for this difficulty with charge changing mutations, as well as finding that FEP+ relative free energies tended to be more positive than experiment.^17^ They further elaborate on FEP+’s difficulty in accurately predicting the effects of Glu, Lys, and Arg mutations due to finite sampling error of sidechain electrostatic interactions in the unfolding reaction.

Another key result of our work is the success of a simple hydrophobic transfer model in predicting the effects of mutations on folding stability. Aside from the seminal work of Eisenberg et al. in characterizing hydrophobic moments of proteins, the key ingredient in this approach appears to be the use of simulated ensembles to accurately capture the solvent accessibility of unfolded protein states. Future work should explore the application of this method to a wider range of proteins, to better gauge its overall accuracy. Modification of the hydrophobic transfer model to include residue-wise secondary structure propensities might further improve its accuracy.

## Conclusion

This work examined the destabilizing effects of ten oncogenic mutations found in the human FOXO1 FKH domain, as seen in DLBCL patients and cell lines. Mutations at L183 greatly reduce FKH stability, and also reduce binding to canonical DNA sequences IRE and DBE2, suggesting that the decrease in folding stability is responsible for the loss of function. To better understand the mechanism of FOXO1 FKH folding and how mutations perturb it, alchemical FEP calculations and massively parallel molecular simulations were performed, enabling the construction of a Markov model that predicts folding pathways, rates, and a mechanism in which the formation of helix α3 is the final step. While FEP overestimates the magnitude of destabilization, a simple hydrophobic transfer model, used in conjunction with simulation-based estimates of solvent-accessible surface area, was shown to quantitatively predict changes in folding free energies from mutations, with high accuracy. These results make a strong case that more realistic, atomistic modeling of unfolded states is highly useful for studying the destabilizing effects of mutations. These observations, together with corresponding disease-associated mutations such as in FOXC1 and FOXG1,^63,64^ support further experimental and computational studies of the folding mechanism of FOX domains.

## Materials and Methods

### Sample Preparation

The FOXO1 Forkhead DNA-binding domain (FKH) was expressed and purified as previously described, with minor modifications.^68^ A codon-optimized fragment encoding NdeI-TEV-FKH (151–265)-XhoI was subcloned into pET28. The construct was transformed into *E. coli* BL21 (DE3), grown in 2xYT medium with 30 μg/ml kanamycin. Protein expression was induced by 0.5 mM IPTG at 21 °C for 8–16 h. Cells were resuspended in 500 mM NaCl, 5 mM beta ME, 20 mM Imidazole, 10% glycerol, disrupted by sonication, clarified by centrifugation and applied to a HisTrap column (GE Healthcare). Protein was eluted with a 0–500 mM imidazole gradient.

His-tagged FKH was desalted into 200-300 mM NaCl, 20 mM HEPES pH 7.5, 0.5 mM EDTA, 1 mM DTT, 5-10% w/v glycerol, cut overnight with His-tagged TEV (1:40 w:w) at 10 °C, followed by passage over Ni-NTA (Qiagen) gravity column to remove TEV and uncut protein. Cleaved FKH protein was exchanged into 50-300 mM NaCl, 20 mM HEPES pH 7.5, 10% w/v glycerol and applied to a MonoS or Capto S column (GE Healthcare). Protein was eluted with a gradient to 1.0 M NaCl. For storage, FKH was concentrated, glycerol added to 25% w/v, flash-frozen in liquid N_2_ and stored at -80 °C.

Biotinylated 32bp DNA oligos were synthesized (Eurofins) containing canonical FKH binding motifs IRE, DBE1 and DBE2 (See Table 1). Complementary oligos were dissolved at 100 μM each in 10 mM Tris-HCI, pH 7.5,100 mM KCl and 1 mM EDTA and annealed in a PCR block with temperature gradient from 80°C to 20 °C over 60 min. Annealed dsDNA was purified by FPLC on a MonoQ 5/50 column (GE Healthcare) with 20 ml linear gradient 0.1–1.1 M KCl. Purified dsDNA was dialyzed against 20 mM Tris-HCl pH 8.0, 50 mM KCl, 5% w/v glycerol, 2 mM DTT, 0.2 mM EDTA, 2 mM MgCl_2_.

### Thermal Shift Assay (TSA)

FKH protein was exchanged into 20 mM Tris-HCl pH 8.0, 50 mM KCl, 5% w/v glycerol, 2 mM DTT, 0.2 mM EDTA, 2 mM MgCl_2_, at 5 μM concentration and 2X Sypro Orange (Thermo Fisher). Aliquots of 50 μl were loaded into PCR strips and analyzed on a StepOne Plus™ qPCR instrument (Thermofisher) with temperature gradient 25–95 °C ramp, 1 °C increment, 1 min hold/increment. Data from the ROX channel was baseline corrected and fit to a 2-state model in Matlab.

### Differential Scanning Calorimetry (DSC)

FKH protein was prepared in Tris-EDTA at 1 mg/ml (∼7.5 μM). Samples were analyzed in a PEAQ-DSC instrument (Malvern) with temperature gradient 25–90 °C. Baseline corrected, buffer-subtracted data were analyzed by a non two-state model.

### Electrophoretic Mobility Shift Assay (EMSA)

FKH at 5–1000 nM in 10 mM Tris-HCl pH 7.5, 50 mM KCl, 5% w/v glycerol, 1 mM DTT, 0.2 mM EDTA, 1 mM MgCl_2_ was incubated with 1 nM biotinylated dsDNA in presence of 1 ng/μl poly-dI/dC for 30 min at room temperature. Samples were separated on 6% 0.5X TBE polyacrylamide gels and analyzed using LightShift® Chemiluminescent EMSA Kit (ThermoFisher).

### Free Energy Perturbation (FEP) Calculations

To model unfolded-state conformations, eight FKH tripeptides were prepared using UCSF Chimera 1.13.1. Alchemical topologies for wild type (wt, A state) and mutant (B state) were prepared using pmx.^19,69^ The tripeptide sequences originating from the FOXO1 FKH domain are T(L183R)S, K(S152R)S, S(S153R)S Y(A166G)D, Y(A166V)D, L(T182M)L, N(S205T)S, and T(L183P)S. Each tripeptide was solvated with TIP3P water in cubic periodic boxes ranging in volume from (3.15 nm)^3^ to (3.54 nm)^3^. Neutralizing K^+^ and Cl^-^ counterions were added at 50 mM, consistent with concentrations used in the TSA and EMSA assays. For charge-changing mutations, (L183R and S153R), a potassium ion was chosen to have charge +1 in the A state, and neutral charge in the B state.

The folded-state model of FOXO1 FKH(151-249)/Mg complex was adapted from PDB 3COA.^6^ Protonation states at pH 7.5 were determined using H++ ^70^ and the AMBER ff14SB force field was used to build the topology.^38^ The calcium ion in 3COA was replaced with magnesium and the MCPB algorithm^71^ was used to build three coordination bonds with the carbonyl oxygens of Leu217, His220, and Phe223. Alchemical topologies were prepared using *pmx*,^69^ with the assistance of in-house code (https://github.com/leiqian-temple/AlchemFEP_FOXO1). The final dual topology used the AMBER 14sbmut forcefield available in *pmx*. Each system was solvated in a cubic periodic boxes ranging in volume from (7.16 nm)^3^ to (7.25 nm)^3^, with TIP3P waters and neutralizing K^+^ and Cl^-^ counterions added at 50 mM. Charge-changing mutations were addressed using the same strategy as the unfolded-state calculations. In addition, six FOXO1 FKH(151-249) conformations corresponding to folding intermediates were taken from the MSM (see <u>Markov Model Construction and Analysis</u> below) and used to prepare alchemical FEP models using a similar process, but with the magnesium-related MCPB steps omitted.

Minimization, equilibration, and production molecular dynamics for the FEP calculations were performed on Temple University’s Owlsnest HPC cluster using the GROMACS 2016.3 simulation package.^72^ Energy minimization was performed until all forces were less than 1000 kJ mol^-1^ nm^-1^. For equilibration and production, stochastic (Langevin) integration was performed with a time step of 1 fs and friction coefficient 1 ps^−1^. PME electrostatics were used with fourier spacing 0.12 and PME order 4. Long-range dispersion correction was used. A total of 30 alchemical intermediates were used to interpolate from the A topology to B topology, using potential energy function *U* = (1-λ)*U*_A_ + λ*U*_B_, where 0 ≤ λ ≤ 1 is the interpolation parameter. The λ values used were (0.000, 0.002, 0.004, 0.006, 0.008, 0.01, 0.02, 0.03, 0.04, 0.05, 0.10, 0.15, 0.20, 0.25, 0.30, 0.35, 0.40, 0.45, 0.50, 0.55, 0.60, 0.65,0.70, 0.75, 0.80, 0.85, 0.90, 0.95, 0.975, 1.00). For each alchemical intermediate, equilibration at 300 K was performed in the NVT ensemble for 100 ps, and then in the NPT ensemble at 1 bar for 100 ps using the Parrinello-Rahman with compressibility constant 4.5 × 10^−5^ and time constant 0.5 ps. Production runs of each model were performed for 8 ns in the NPT ensemble. For each simulation performed using a given value λ_*i*_, the values of ∂*U*/∂λ and Δ*U*_*ij*_ = *U*(λ_*j*_) – *U*(λ_*i*_) were written to file every 0.5 ps, for all intermediates *j*. Free energy estimation was performed with the Multistate Bennett Acceptance Ratio (MBAR) method,^73^ using the alchemical-analysis^74^ and pymbar^75^ Python packages. Time-reversed convergence plots for Δ*G* estimates were used to identify and discard the non-equilibrated regions of the trajectories (<u>Figure S6</u>). These results show that most simulations meet conditions of equilibrium and convergence in the last 4 ns, with the possible exception of folded-state S152R. Therefore, we removed the first 4 ns of each 8-ns trajectory before analysis. The overlap matrix, whose elements *O*_*ij*_ quantifies the overlap between the distributions of Δ*U*_*ij*_,^74^ shows sufficient overlap for accurate free energy estimates (<u>Figure S7</u>).

### Massively Parallel Folding Simulations

The model of FOXO1 was adapted from PDB 3CO6. ^6^ Protonation states at pH 7.5 were determined using H++,^70^ and the AMBER14SB force field was used to build the topology.^38^ The protein was placed in a cubic periodic box of volume (8.2698 nm)^3^ and solvated with TIP3P waters,^39^ with Joung-Cheatham neutralizing Na^+^ and Cl^-^ counterions added at 0.1 M,^76,77^ resulting in a system of ∼64500 atoms. Molecular dynamics simulations were performed using the GPU-accelerated OpenMM (CUDA platform), using stochastic (Langevin) integration with a 2 fs time step, friction coefficient of 1 ps^-1^ and PME electrostatics with cutoff 1.0 nm and tolerance 0.005.^37,78^ To generate a range of folded and unfolded conformations, several 60-ns NPT simulations were performed at 300 K, 400 K, 450 K, and 498 K, respectively. Conformational clustering was performed using a *k*-centers algorithm to identify representative conformations.^79^ Production runs were performed at 375 K on the Folding@home distributed computing platform,^32^ with all coordinates saved every 0.5 ns. Five hundred trajectories each with randomized initial velocities were initiated from 20 different starting structures, resulting in 10,000 total independent simulations. An aggregate of ∼6.8 ms of simulation data was collected.

### Markov Model Construction and Analysis

Using PyEMMA 2.5.7,^80^ all 9927 obtained trajectories were featurized using the pairwise distance for every other C*α* in residues 5 to 95 for a total of 1035 features. Dimensionality reduction was performed using time-lagged Independent Component Analysis (TICA) with a lag-time τ_TICA_ = 2.5 ns. A *k*-means algorithm was used to conformationally cluster in the low-dimensional TICA projections to define MSM microstates. The VAMP2 score was used to assess models constructed with different numbers of microstates (<u>Figure S8</u>) and different low-rank TICA projections. Based on this score, and a general desire for computational tractability, we selected an MSM with 1000 microstates, using the four largest TICA components. MSM implied timescales were calculated as *t*_*i*_ = -τ/ln µ_*i*_, where µ_*i*_ are the largest nonstationary eigenvalues of the microstate transition matrix. The slowest timescale was stable after 5.0 ns. A Bayesian MSM with a lag time of τ = 5.0 ns was constructed using these microstate definitions.^42^ The microstates were assigned to macrostates (U, I and N) using the PCCA+ algorithm.^43,44^ This coarse-grained three-state model passed the Chapman-Kolmogorov test (<u>Figure S9</u>). Mean first passage times from macrostates A to B were calculated using PyEMMA, as the self-consistent expectation value *E*_A_ of the time *T*_B_ to reach a state in B from a state in A, *E*_A_[*T*_B_] = Σ_*a*∈A_ (π_*a*_ *E*_*a*_[*T*_B_]/Σ_*z*∈*A*_ π_*z*_), where π_*a*_ are the equilibrium populations of state *a*.

All trajectory analysis was performed through a series of in-house Python 3.7.6 scripts utilizing MDTraj 1.9.5 and Numpy 1.18.1 libraries, unless otherwise stated. Visualization was performed using Matplotlib 3.3.4 and VMD 1.9.3. Per-residue secondary structure probabilities for each macrostate were calculated via the DSSP algorithm, by comparing the frequency of trajectory snapshot DSSP assignments to the DSSP assignments of the crystal structure. Secondary structural elements were defined by the following residue ranges: *α*1 (165–175), *α*2 (183–193), *α*T (195–197), *α*3 (203–219), *β*4 (223–226), *β*5 (236–239).

The frequency of native contacts for each macrostate was calculated using the sigmoidal function (1+exp(β(*d*_*ij*_ - λ*d*_0_))^−1^ to count contacts, where *d*_*ij*_ are the distances between atoms *i* and *j*, β = 50 nm^-1^, λ = 1.8, *d*_0_ = 0.45 nm, as described by Best et al. (2013).^46^ We adapted this method to coarse-grain the contact description as residue-wise rather than atomic, such that if an atomic contact is present between two residues, those residues are considered in contact. The contact frequencies for each macrostate were calculated as a population-weighted average across all microstates assigned to the macrostate.

The Shrake-Rupley algorithm as implemented in MDTraj was used to calculate the solvent-accessible surface area (SASA) for 20 trajectory samples chosen at random from each microstate.^49^ The average residue-wise SASA for each macrostate was then calculated as a population-weighted average across all microstates assigned to the macrostate, where the microstate populations come from the MSM.

### A hydrophobic transfer model of ΔΔ*G*_fold_

To make quantitative predictions about the about the effects of mutations on protein stability, we developed a transfer free energy model of ΔΔ*G*_fold_ based on the work of Eisenberg et al. on environmental hydrophobicity.^50,51^ In this model, the free energy of a protein is given in terms of residue hydrophobicities *H*(*r*_*i*_) for each residue *r*_*i*_ at sequence position *i*, and the environmental hydrophobicity of each residue, *M*_*i*_.

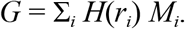

The hydrophobicities are per-residue free energies, representing the free energy of transfer from a hydrophobic to a hydrophilic phase (specifically, the “consensus hydrophobicities” compiled by Eisenberg et al.^81^). The environmental hydrophobicities are assumed to depend on solvent exposure, such that *M*_*i*_ = *A*(*r*_*i*_)/*A*_0_(*r*_*i*_) – ½, where *A*(*r*_*i*_) is the conformation-dependent solvent exposure of residue *r*_*i*_, and *A*_0_(*r*_*i*_) is the maximum possible solvent exposure of residue *r*_*i*_. In this way, the values of *M*_*i*_ range from a fully aqueous environment of *M* = +½, to an environment of full burial in the interior of a protein, *M* = -½.

According to this model, the free energy of folding is

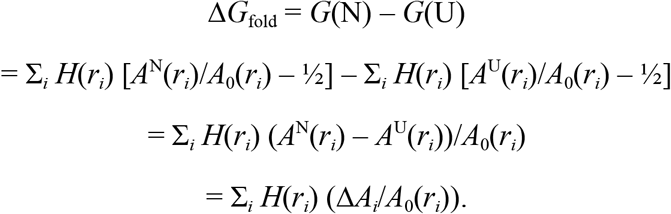

Here, Δ*A*_*i*_ denotes the change in the solvent exposure of residue *i* upon folding. For a single-point mutation at position *i* from residue *r*_*i*_ (wild type) to *s*_*i*_ (mutant), the change in the free energy of folding, ΔΔ*G*_fold_ (U→N), depends *only* on residue *i*,

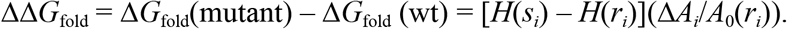

Thus, the model posits that ΔΔ*G*_fold_ is the change in the hydrophobicity upon mutation Δ*H*(*r*_*i*_ → *s*_*i*_), multiplied by the fractional change Δ*A*_*i*_/*A*_0_(*r*_*i*_) in the solvent exposure upon folding. In practice, we obtain values of *A*_0_(*r*_*i*_) from the normalized maximal accessible surface area scale of Tien et al.^53^ Consensus hydrophobicities and *A*_0_ parameters used in our model are listed in <u>Table S2</u>.

### Estimation of folding free energies from Δ*T*_*m*_

To estimate the change in the folding free energy ΔΔ*G*_fold_ from the experimental values of Δ*T*_m_, we assume the standard model of protein stability temperature-dependence,^20^

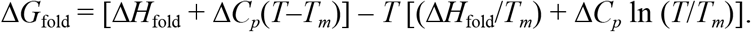

We use the work of Robertson and Murphy^21^ to estimate Δ*H*_fold_ and Δ*C*_*p*_ from the number of residues *N*,

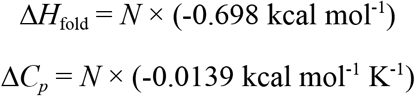

Using these values with *N*=100, we estimate ΔΔ*G*_fold_ as

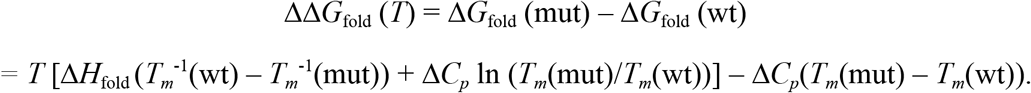

Results in this manuscript are reported at *T* = 298.15 K.

## Acknowledgments

We thank the participants of Folding@home, without whom this work would not be possible. This work was supported in part by the National Institute of General Medical Sciences (1R01GM114358) of the National Institutes of Health to RHGB. DN and VAV were supported by NIH 1R01GM123296. Temple HPC resources were supported by NSF CNS-162506 and US Army Research Laboratory W911NF-16-2-0189. The CB2RR Computing resource is supported by NIH S10-OD020095.

## Data Availablity

All simulation trajectory data and analysis scripts are available via the Open Source Framework at https://osf.io/t7h5b/.

## Supporting Information

Supporting Tables S1 and S2, and Figures S1–S9.

Supporting Movies M1–M8.

