## Supporting Information for "Oncogenic mutations in the DNA-binding domain of FOXO1 disrupt folding: quantitative insights from experiments and molecular simulations"

### Supporting Tables

**Table S1.**

Mean first passage times (MFPTs) between macrostates.

| MFPT (μs) | to U | to I | to N |
| --- | --- | --- | --- |
| from U | - | 9.4 | 20.7 |
| from I | 6.4 | - | 6.9 |
| from N | 15.5 | 5.9 | - |

**Table S2.**
Hydrophobic transfer (HT) model parameters for each amino acid. Normalized maximal accessible surface areas $A_0$ are taken from Tien et al.[53] Consensus hydrophobicities are taken from Eisenberg et al.[81]

| Residue | $A_0$ ($nm^2$) | Consensus hydrophobicities (kcal/mol) |
| --- | --- | --- |
| Glycine | 0.97 | 0.48 |
| Alanine | 1.21 | 0.62 |
| Serine | 1.43 | -0.18 |
| Arginine | 2.65 | -2.53 |
| Proline | 1.54 | 0.12 |
| Threonine | 1.63 | -0.05 |
| Methionine | 2.03 | 0.64 |
| Valine | 1.65 | 1.08 |
| Leucine | 1.91 | 1.06 |

Supplemental Figures

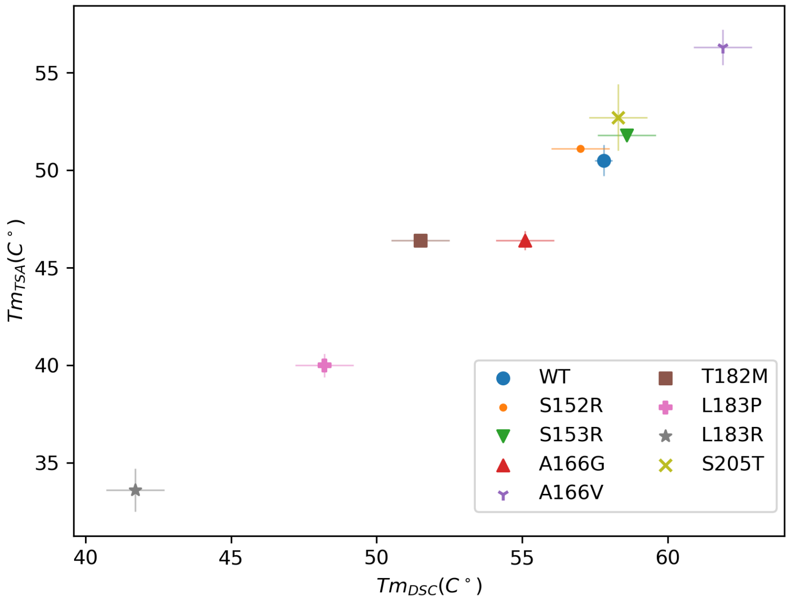


**Figure S1.**

Comparison of melting temperatures $T_m$ measured in thermal shift assays (TSA) versus those measured using differential scanning calorimetry DSC. Uncertainties in DSC $T_m$ measurements for mutants are estimated to be ±1 C.

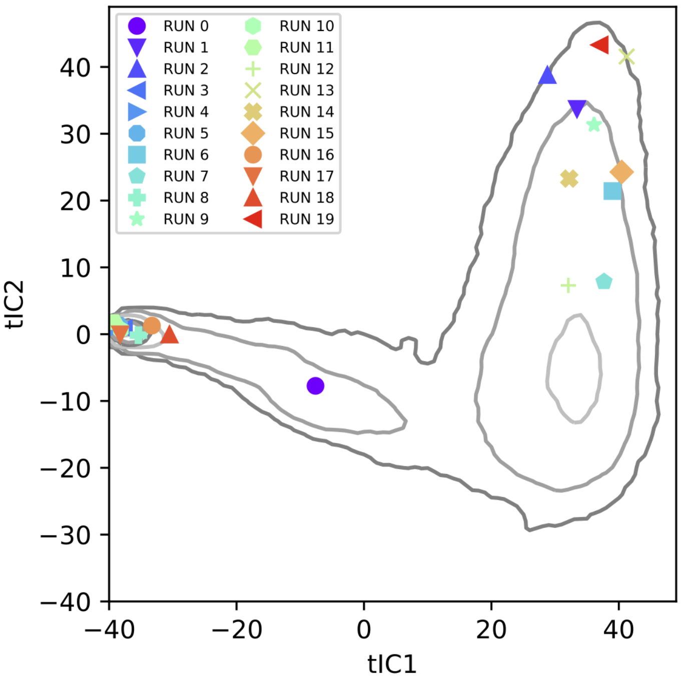


**Figure S2.**

The locations of the twenty conformations used to initiate trajectories, projected onto the (tIC1, tIC2) free energy landscape.

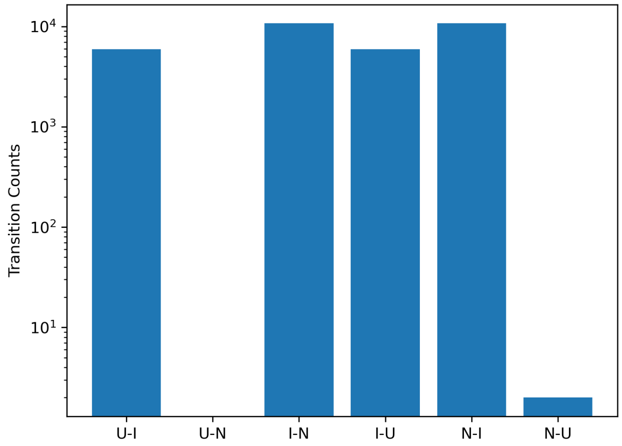


**Figure S3.**

Transition counts observed between macrostates U, I and N (at lag time 5.0 ns) in the simulation trajectory data.

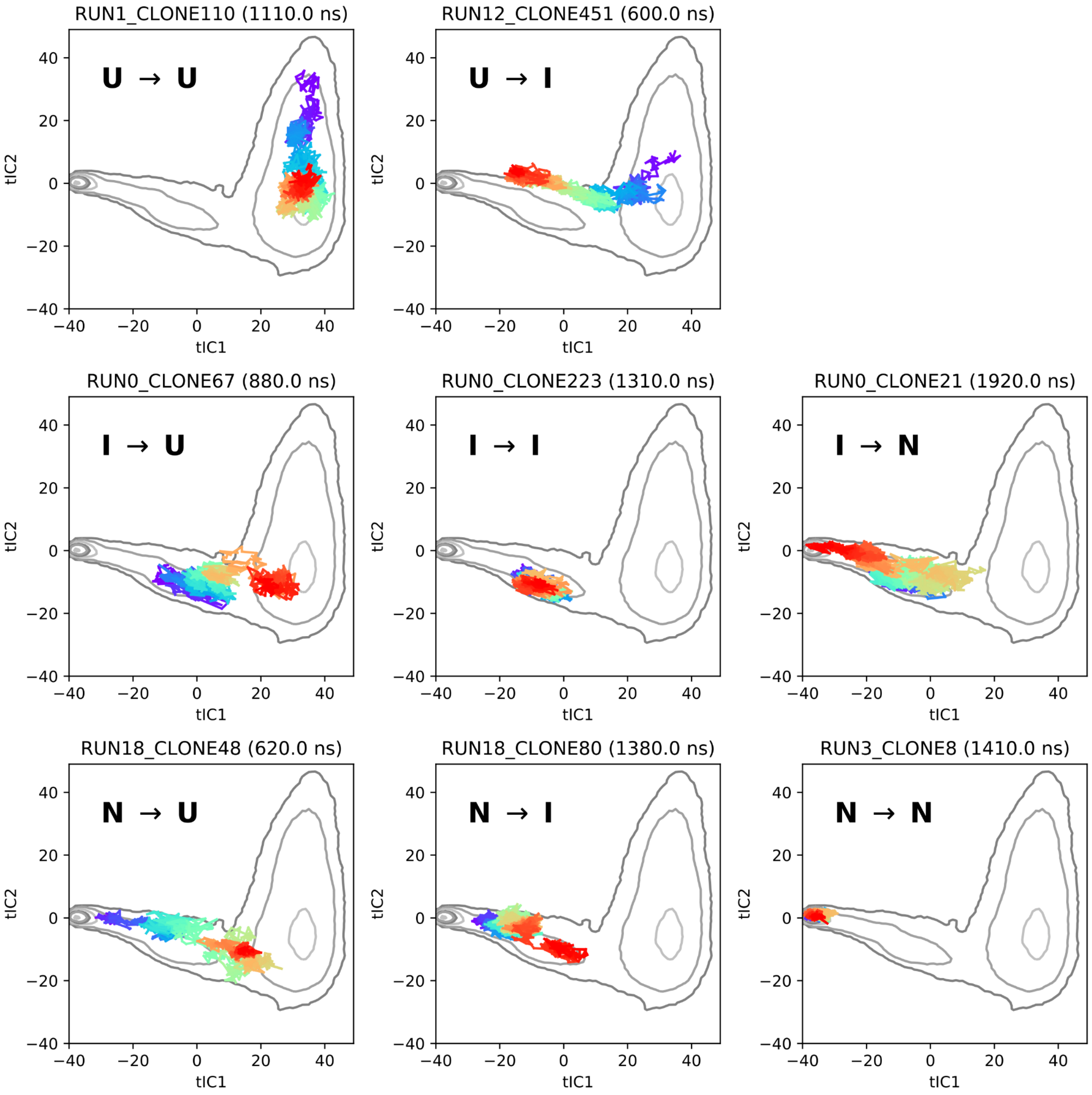


**Figure S4.**

Examples of trajectories involved in observed macrostate transitions. Traces are plotted on the TICA free energy landscape, with the color denoting the direction of time (violet to red denotes the start to finish). Animated visualizations of each of these trajectories are included in Supplemental Movies M1–M8.

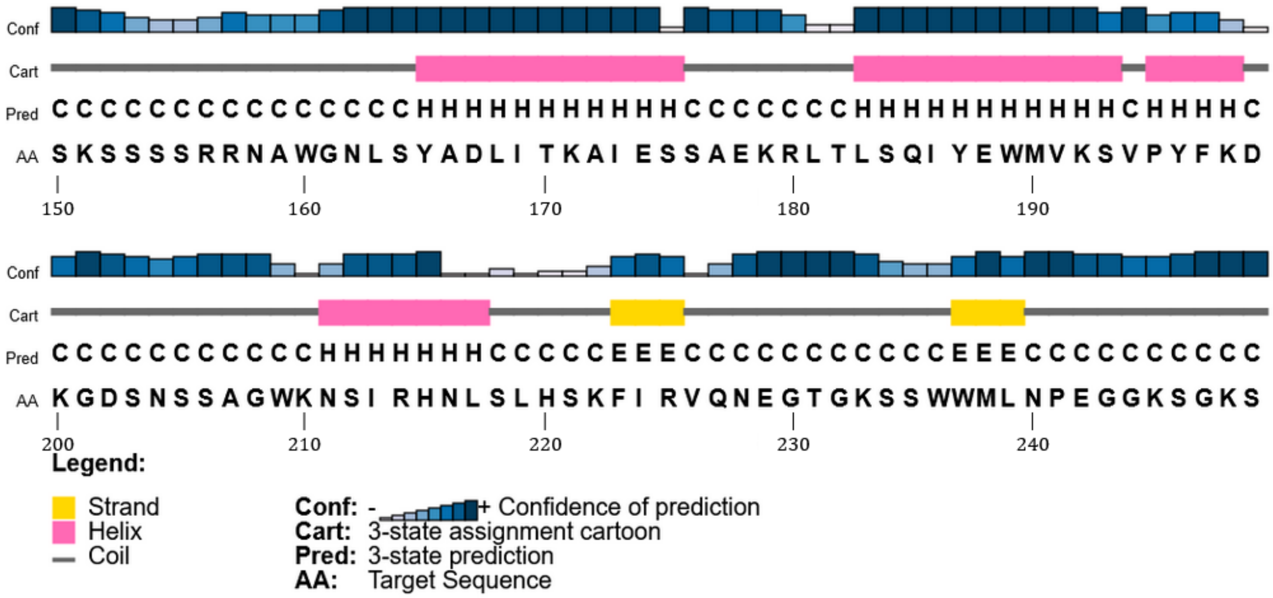


**Figure S5.**

PSIPRED (http://bioinf.cs.ucl.ac.uk/psipred/) predictions of FOXO1 secondary structure.

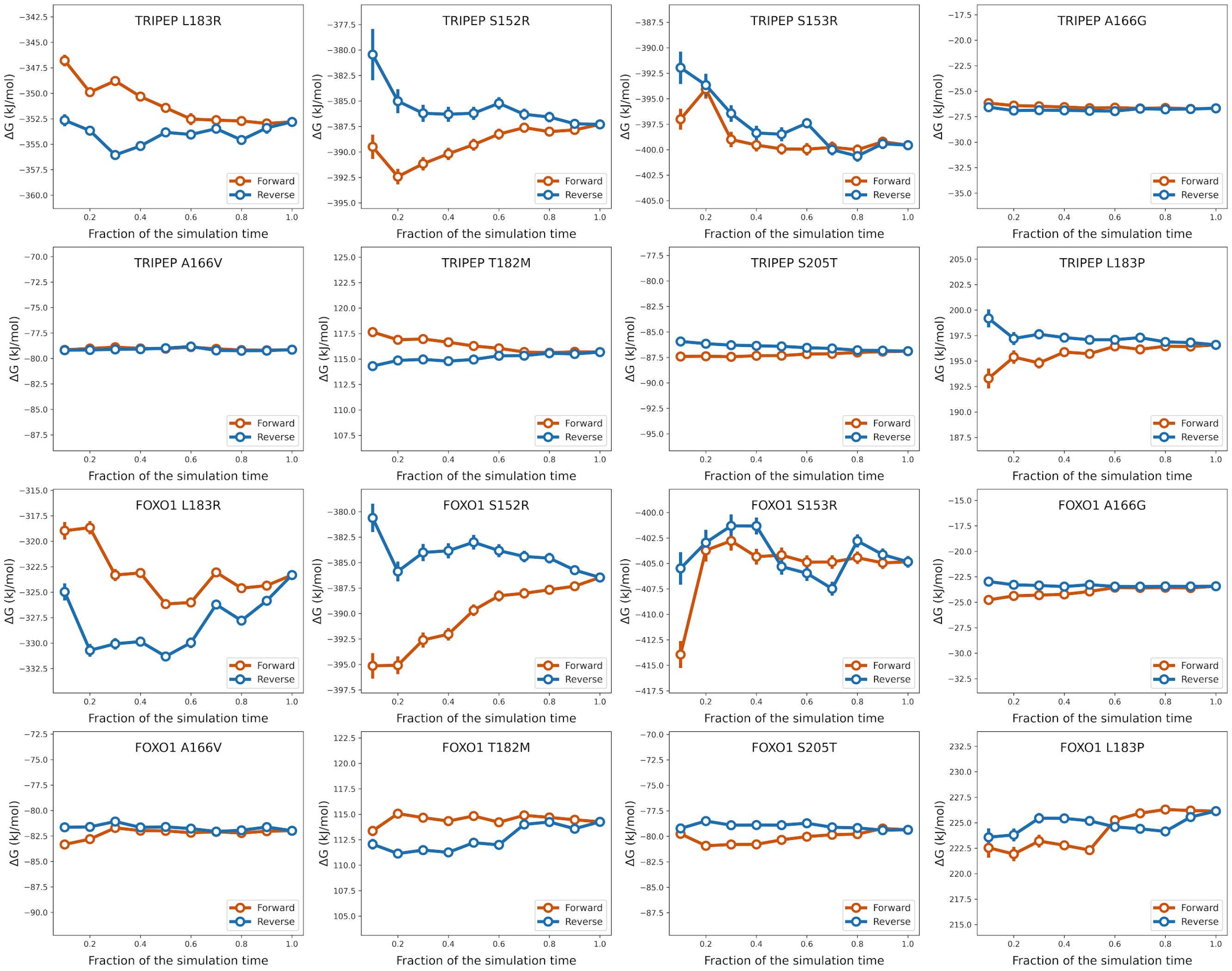


**Figure S6.**

Time-reversed convergence plots for $\Delta G$ estimates were used to identify and discard non-equilibrated regions of the trajectories. In this analysis, two calculations are compared in which increasing amounts of data from a 8-ns trajectory are given to the MBAR free energy estimator; one from a time-forward trajectory and the other from a time-reversed trajectory. When all the data is included, the two calculations yield the same estimate; disagreement of the estimates when only part of the data is included may indicate convergence problems. The results show that most of the FEP simulations are sufficiently converged by the last 4 ns, with the possible exception of folded-state S152R. Therefore, we report FEP estimates computed from only the last 4 ns of each 8-ns trajectory.

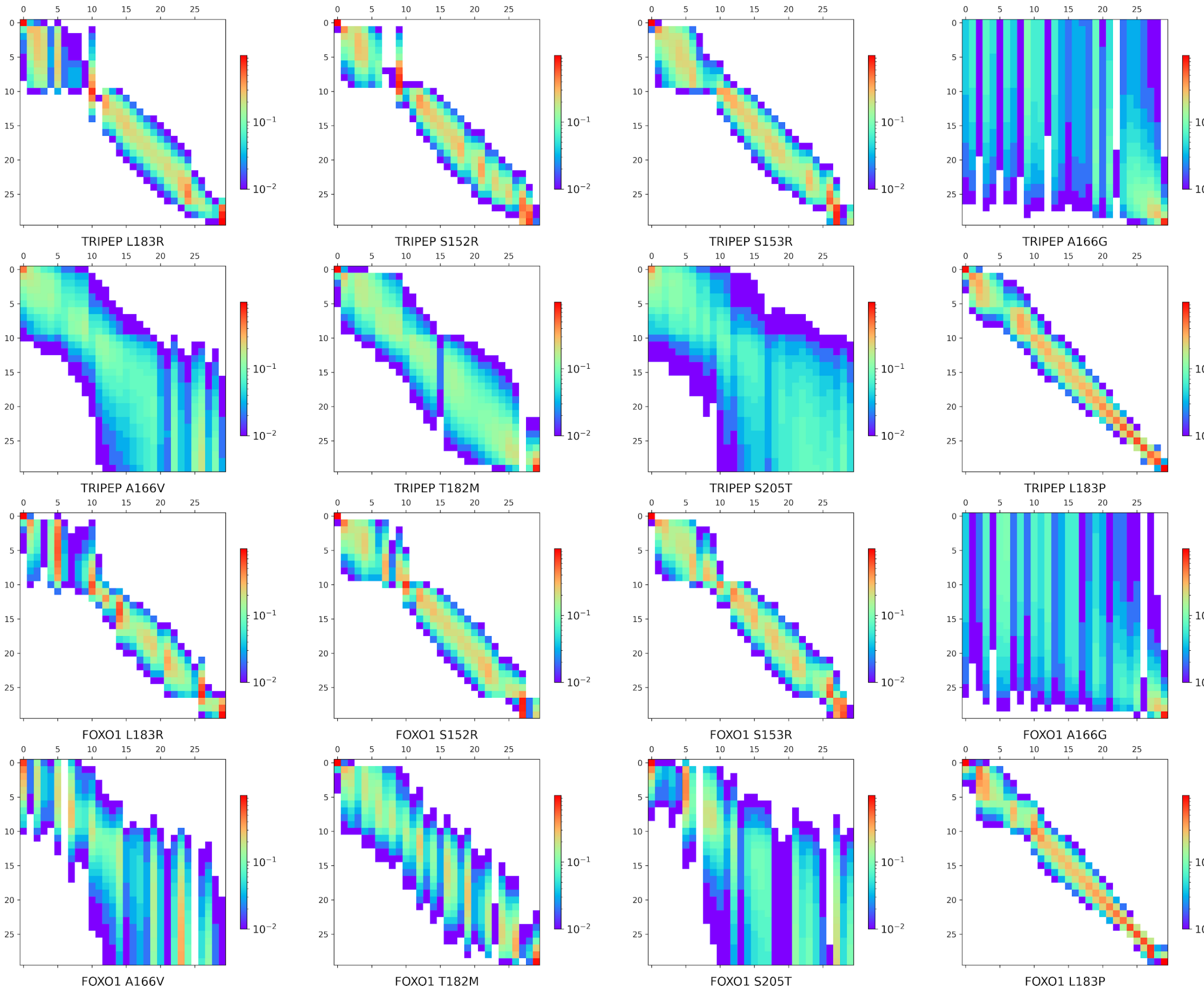


**Figure S7.**

The overlap matrix, whose elements $O_{ij}$ quantify the overlap between the distributions of $\Delta U_{ij} = U(\lambda_j) - U(\lambda_i)$ for alchemical intermediate indices $i$ and $j$ (see Klimovich et al. 2015), is shown for each FEP calculation.

A

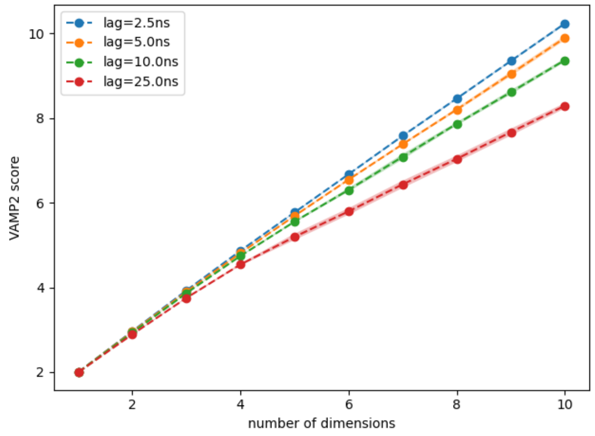


B

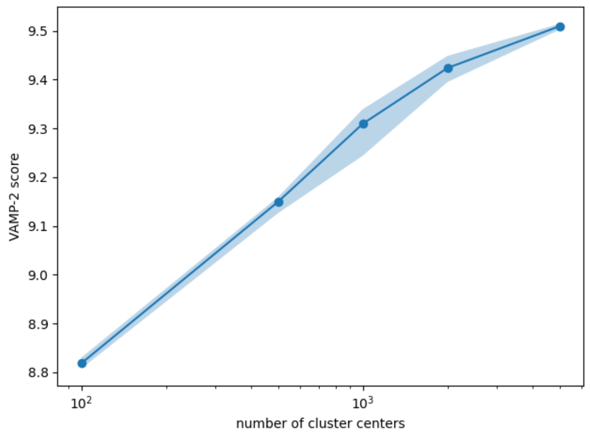


**Figure S8.**

(A) To choose an appropriate TICA lag time and number of TICA dimensions, VAMP2 scores were computed for different microstate MSMs constructed using various numbers of TICA dimensions and TICA lag times: $\tau_{\mathrm{TICA}}$ = 2.5 ns (blue), 5.0 ns (orange), 10.0 ns (green), and 25.0 ns (red). All models used alpha-carbon distance features, and 1000 *k*-means cluster centers. Based on these results, we chose to construct MSMs of FOXO1 using a TICA lag time of $\tau_{\mathrm{TICA}}$ = 2.5 ns, and four TICA dimensions. (B) To determine an appropriate number of cluster centers, VAMP2 scores were computed for microstate MSMs constructed using different numbers of cluster centers. All calculations were performed using an MSM lag time of $\tau_{\mathrm{MSM}}$ = 5.0 ns and TICA lag time of $\tau_{\mathrm{TICA}}$ = 2.5 ns.

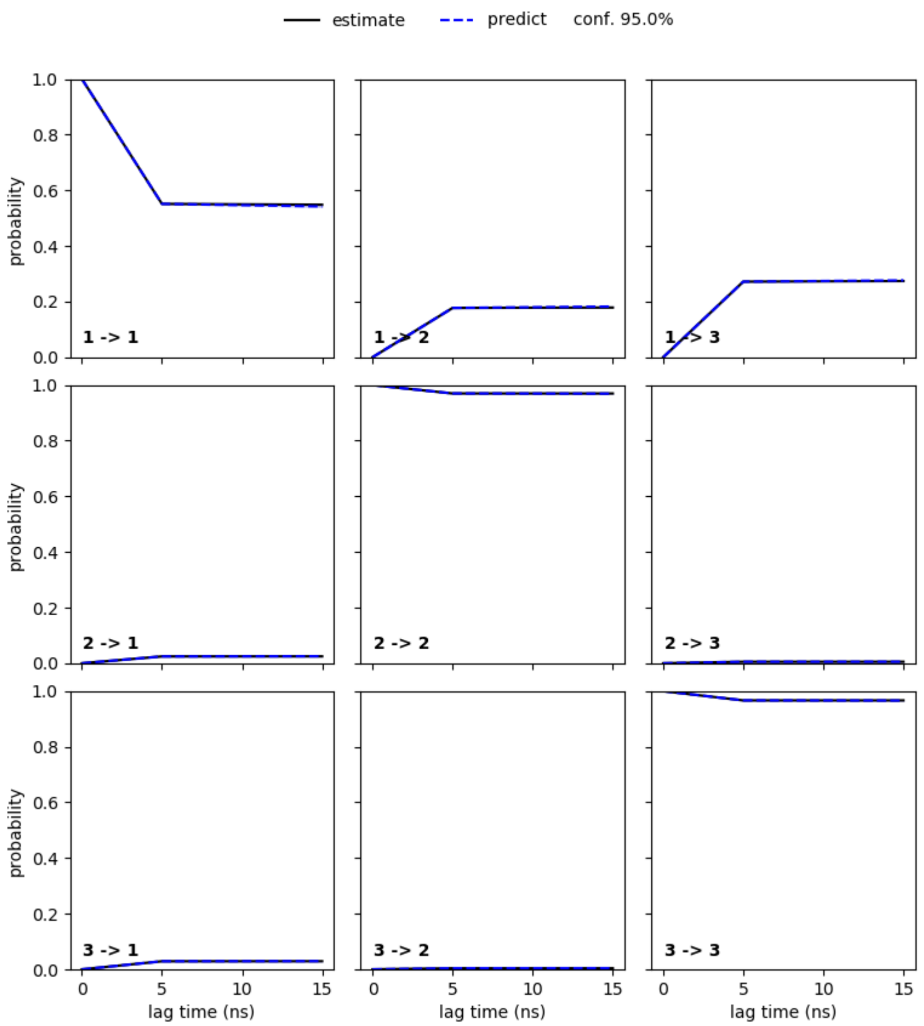


**Figure S9.**

The Chapman-Kolmogorov test is used to check whether the macrostate MSM is sufficiently Markovian. In this test, empirical estimates of the 3 × 3 macrostate transition probability matrix $\mathbf{T}^{(n\tau)}$ constructed at lag times $n\tau$, $n$ = 1, 2, 3… (solid lines) are compared to predictions $[\mathbf{T}^{(\tau)}]^n$ (dashed lines). Each row of panels labeled "$i \rightarrow j$" shows the time evolution of populations for macrostate $j$ where the initial population is placed entirely in macrostate $i$. Here, $i$=1 is macrostate I, $i$=2 is macrostate N, and $i$=3 is macrostate U. The results show that the two models (empirical vs. prediction) are virtually indistinguishable, indicating the dynamics is Markovian.

Supplemental Movies

**Movie M1**. Visualization of a U→U trajectory (1110 ns, RUN 1, CLONE 10).

**Movie M2**. Visualization of a U→I trajectory (600 ns, RUN 12, CLONE 451).

**Movie M3**. Visualization of a I→U trajectory (880 ns, RUN 0, CLONE 67).

**Movie M4**. Visualization of a I→I trajectory (1310 ns, RUN 0, CLONE 223).

**Movie M5**. Visualization of a I→N trajectory (1920 ns, RUN 0, CLONE 21).

**Movie M6**. Visualization of a N→U trajectory (620 ns, RUN 18, CLONE 48).

**Movie M7**. Visualization of a N→I trajectory (1380 ns, RUN 18, CLONE 80).

**Movie M8**. Visualization of a N→N trajectory (1410 ns, RUN 3, CLONE 8).
